# Genome writing to dissect consequences of SVA retrotransposon disease X-Linked Dystonia Parkinsonism

**DOI:** 10.1101/2025.10.07.680816

**Authors:** Weimin Zhang, Yu Zhao, Priya Prakash, Heather L. Appleby, Kelly Barriball, Simona Capponi, Qingwen Jiang, Aleksandra M. Wudzinska, Christine A. Vaine, Gwen Ellis, Neha Rahman, Stefan Markovic, Orin Mishkit, Kerry C. Limberg, Matthew T. Maurano, Youssef Z. Wadghiri, Sang Yong Kim, H. T. Marc Timmers, D. Cristopher Bragg, Shane A. Liddelow, Ran Brosh, Jef D. Boeke

## Abstract

Human retrotransposon insertions are often associated with diseases. In the case of the neurodegenerative X-Linked Dystonia-Parkinsonism disease, a human-specific SINE-VNTR-*Alu* subfamily F retrotransposon was inserted in intron 32 of the *TAF1* gene. Here, we genomically rewrote a portion of the mouse *Taf1* allele with the corresponding 78-kb XDP patient derived *TAF1* allele. In mESCs, the presence of the intronic SVAs—rather than the hybrid gene structure—reduces hy*TAF1* levels. This leads to transcriptional downregulation of genes with TATA box enriched in their promoters and triggering apoptosis. Chromatin and transcriptome profiling revealed that intronic SVAs are actively transcribed, forming barriers that likely impede transcription elongation. In mice, neuronal lineage *TAF1* humanization resulted lethality of male progeny within two months. XDP male mice had severe atrophy centered on the striatum—the same affected brain region in XDP patients. Lastly, CRISPRa-mediated activation of hy*TAF1* restored mESC viability, suggesting boosting *TAF1* transcription as a therapeutic approach.

## Introduction

SINE-VNTR-*Alu* (SVA) elements are a class of hominoid-specific, non-autonomous retrotransposons that are active in the human genome^1,2^. A canonical SVA element comprises a CCCTCT hexamer repeat, two antisense *Alu*-like sequences, a variable number of tandem repeat (VNTR) region, and a SINE-R domain followed by a poly A-tail^1^. Although SVAs are non-autonomous, they are mobilized through target-primed reverse transcription (TPRT), a mechanism dependent on LINE-1 reverse transcriptase^3,4^. Based on phylogenetic sequence analyses of the SVA and 5′ transduced sequences, SVAs are categorized into seven subfamilies, SVA_A to SVA_F, plus SVA_F1^5,6^. Similar to other transposable elements (TEs), the *cis*-regulatory activities of SVAs are mostly repressed by Krüppel-associated box (KRAB) domain-containing zinc-finger proteins^7^. Specifically, ZNF91 was shown to recruit TRIM28 to its SVA targets and induce local transcriptional silencing by establishing mini-heterochromatin domains^8^.

SVA insertions have been implicated in many human diseases^9–12^ and in the evolution of human pigmentation^13^. Germline SVA retrotransposition could lead to founder-effect genetic diseases, such as the X-Linked Dystonia-Parkinsonism (XDP). XDP is an adult-onset progressive movement disorder with high penetrance. XDP is endemic to the Panay Island in the Philippines and affects descendants of this population worldwide. XDP patients manifest characteristic patterns of retrocollis, oromandibular dystonia and tongue protrusion, accompanied with parkinsonism as the disease progresses^14^. Magnetic resonance imaging (MRI) of patient brains revealed atrophy of the striatum, underlining the neurodegenerative nature of the disease. Genetic studies have identified the causal variant as a SINE-VNTR-*Alu* (SVA) retrotransposon insertion in intron 32 of the *TAF1* gene^15^ (**Fig. 1A**), which encodes the largest subunit of the TFIID complex, a central component of the core transcriptional machinery that recognizes the core promoter of genes for RNA polymerase II-mediated transcription^16,17^.

**Figure 1.**
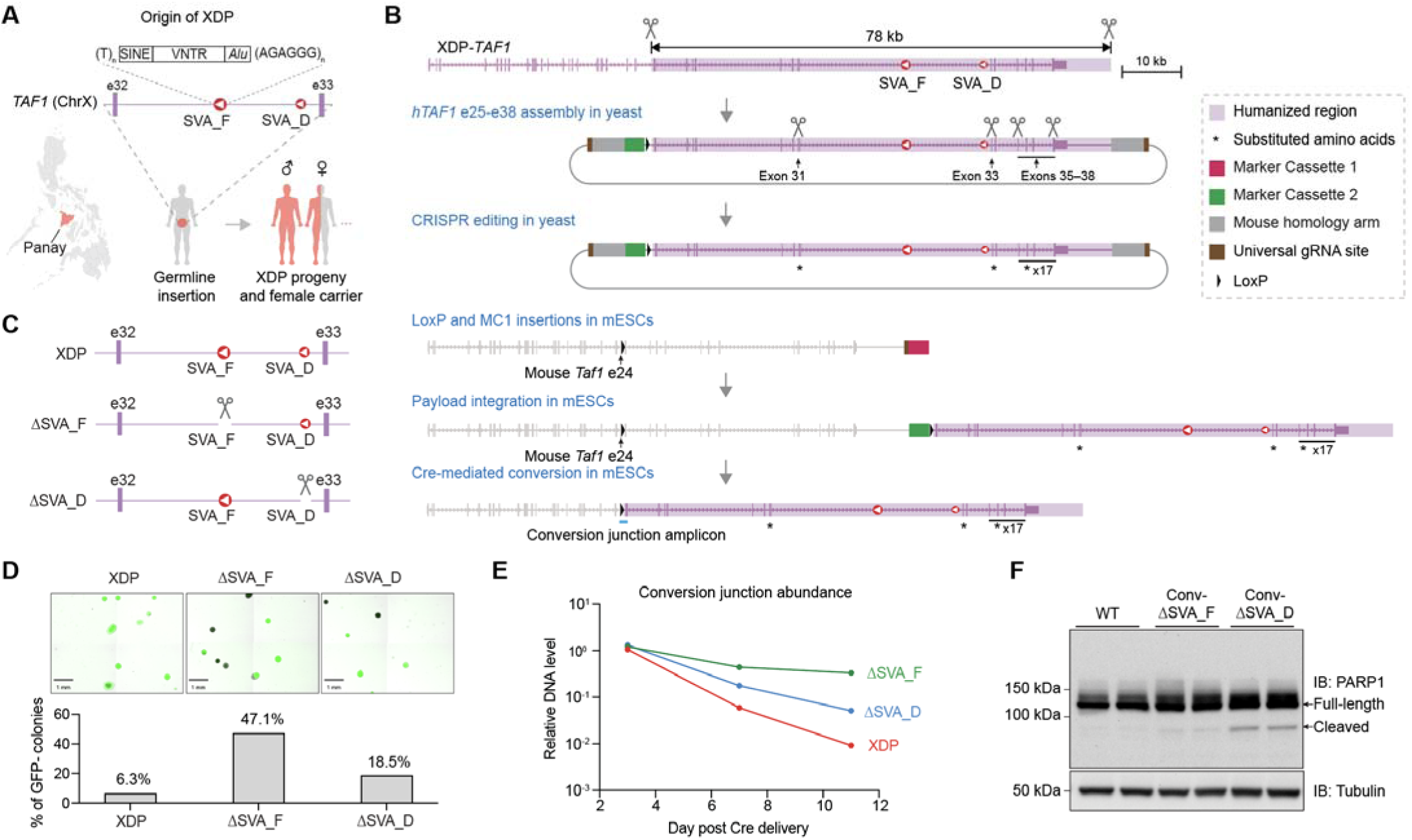
Engineering a conditional partially humanized *TAF1* allele in mESCs. (A) Genetic origin of XDP. The genetic variation of XDP is an SVA_F insertion into intron 32 of the *TAF1* gene in the reverse orientation on chromosome X. An ancestral SVA_D in the same intron is common to all humans. The disease is from a common founder; male progeny (red human cartoon) inheriting the XDP *TAF1* allele manifest disease; while heterozygous (red/grey cartoon) females are typically unaffected or display mild phenotypes. Panay island is highlighted in red, showing the most prevalent region of the disease. e32 and e33, exons 32 and 33 of *TAF1*. (B) Strategy for conditional partial humanization of *Taf1* in mESCs. A genomic fragment of an XDP patient spanning *TAF1* intron 24 to 7.6 kb downstream of the 3’ UTR (hg38: chrX:71398875-71473639) was assembled into a yeast assembly vector (YAV), which harbors flanking universal gRNA sites (brown); two 2.1-kb homology arms targeting a region 6.6 kb downstream of mouse *Taf1* (grey); a GFP containing marker cassette followed by a loxP site (green; see details in **Fig. S1B**). The assembled payload was edited using CRISPR in yeast^25^ to substitute all divergent human TAF1 amino acids to that of mouse TAF1 (substitutions are denoted by asterisks). The payload construct was integrated into a pre-engineered mESC line with a loxP site inserted in intron 24 of *Taf1*, and an mScarlet containing marker cassette 1 (MC1, **Fig. S1D**) inserted 6.6 kb downstream of the mouse *Taf1* 3’ UTR, resulting in a pre-converted hy*TAF1* allele. Cre-mediated excision of mouse exons 25–38 and the GFP containing marker cassette 2 forms the converted-hy*TAF1* allele. (C) CRISPR deletion of either SVA_F or SVA_D from human *TAF1* in yeast. The ΔSVA_F and ΔSVA_D payloads were delivered into mESCs via mSwAP-In. Coordinates of sgRNAs used for SVA_F and SVA_D deletion are listed in **Supplementary Table 2**. (D) Representative images of day 9 post Cre-mediated conversion and quantification of conversion efficiency. GFP-negative colonies indicate successful conversion, scale bar is 1 mm. Bar graph on the bottom summarizes the percentages of GFP negative clones. (E) Quantitative PCR analysis using a novel conversion junction-specific primer pair (oWZ2213 and oWZ2214, amplicon is shown as a blue bar in Fig. 1B). A genomic mouse *Actb* primer pair (oWZ1683 and oWZ1684) was used as an internal reference. (F) Immunoblot of PARP1 in wild-type, converted-ΔSVA_F and converted-ΔSVA_D clones. Full-length (113 kDa) and cleaved (89 kDa) PARP1 are indicated by arrows. α-Tubulin serves as loading control. Two independent mESC clones were analyzed for each genotype.

Modeling exon-disruptive SVA insertions in mice is relatively straightforward, as they typically lead to loss-of-function mutations. In contrast, intronic insertions—such as the one seen in XDP—are more challenging to study, since they often interfere with splicing or regulatory elements. Overcoming this challenge requires faithful humanization of the affected gene and the surrounding genomic context. To enable this, we have developed a large-scale (up to ∼200 kb in a single step) mouse genome rewriting and tailoring method^18^, providing tools to engineer SVA-containing human genes into the mouse genome, and enable faithful modeling of SVA pathologies in mice.

Here, we partially humanized mouse *Taf1* allele with an XDP human patient-derived *TAF1* gene that includes the pathogenic SVA_F insertion and is conditionally expressed following Cre-mediated recombination. The resulting mouse embryonic stem cells (mESCs) and mouse models recapitulate key molecular and pathological hallmarks of XDP and establish a foundation for mechanistic dissection and preclinical therapeutic development. The conditionally humanized *TAF1* mice display symptoms up to 300 times faster than human patients (2 months vs. 50 years), which is likely due to the absence of human specific *ZNF91*^8,19,20^ and perhaps other primate-specific *ZNF* genes from the mouse genome, and possibly because mice mature much faster. The amplified pathologic phenotypes in mice could favor rapid therapeutics development. We show here that introducing CRISPRa reagents activating hy*TAF1* expression in mESCs can alleviate the lethality caused by the XDP-associated SVA_F insertion.

## Results

### Engineering a partially humanized *TAF1* allele in mESCs

To model the effect of the XDP-causing SVA_F insertion in intron 32 of *TAF1* with minimal disruption to the endogenous regulatory architecture of this essential gene, we engineered a conditional partially humanized *Taf1* allele in mESCs. This gene, termed hybrid *TAF1* (hy*TAF1*), comprises mouse *Taf1* exons 1–24 (with intervening introns) and human XDP patient-derived *TAF1* exons 25–38 (with intervening introns) and is only expressed following Cre-mediated recombination, a process we refer to as “conversion”. (**Fig. 1B, Fig. S1A**). The 78-kb human *TAF1* genomic segment (**Fig. 1B** purple region) harbors both the XDP-specific SVA_F (with 34 contiguous hexametric repeats) and an ancestral SVA_D element (common to all humans) in intron 32, as well as 7.6 kb of downstream genomic sequence. To engineer this complex allele, we first cloned the human *TAF1* region into a yeast assembly vector (YAV) that contained mouse *Taf1*-targeting homology arms, flanking sgRNA sites for linearization *in vivo*, and a selectable marker cassette (MC2) followed by a loxP site, enabling integration downstream of mouse *Taf1* via mSwAP-In^18^ (**Fig. S1B**). To mitigate potential cross-species incompatibility, nineteen divergent amino acids in the human TAF1 protein were substituted to match the mouse sequence, yielding a protein sequence identical to wild type mouse TAF1 (**Fig. 1B** asterisks, **Fig. S1C**). Male mESCs were pre-engineered to harbor a loxP site in intron 24 of mouse *Taf1*, and a distinct marker cassette (MC1) 6.6 kb downstream of mouse *Taf1* gene (**Fig. S1D**). Using mSwAP-In^18^, MC1 was then swapped with the human *TAF1* payload, resulting in a “pre-converted” hy*TAF1* allele (**Fig. 1B**). Human *TAF1* DNA copy number was quantified to ensure single copy on chromosome X (**Fig. S1E** and **Methods**). Three independent pre-converted clones were validated by targeted sequencing^21^ (**Fig. S1F**), and the presence of SVA_F was confirmed by PCR as a ∼3 kb amplicon (**Fig. S1G**).

Upon Cre-mediated excision of the endogenous mouse *Taf1* exons 25–3’ UTR region and marker cassette 2, the conversion junction amplicon was readily detected by PCR and sequencing (**Fig. S1H**). However, the converted mESCs expressing the hy*TAF1* gene underwent cell death and failed to form viable clones (**Fig. S1I**). Longitudinal conversion junction PCR quantification of Cre-transfected cells revealed a rapid decline in the amount of the conversion junction amplicon, consistent with rapid elimination of converted mESCs (**Fig. S1J**). To determine which SVA element in intron 32 contributes to this cytotoxicity, we engineered pre-converted mESC lines lacking either SVA_F or SVA_D (hereafter ΔSVA_F, ΔSVA_D, **Fig. 1C**) and subjected them to Cre-mediated conversion. We monitored conversion efficiency by measuring the loss of GFP fluorescence in emerging mESC colonies since GFP is excised along with MC2 and mouse *Taf1* e25–3’ UTR during conversion. Approximately 47.1% of the ΔSVA_F colonies and 18.5% of the ΔSVA_D colonies lost GFP expression, indicating successful conversion. In contrast, only 6.3% of the XDP mESC colonies were GFP negative, but these colonies were markedly smaller than their ΔSVA_F and ΔSVA_D counterparts and were likely composed of apoptotic cells (**Fig. 1D**). These converted XDP colonies failed to expand after manual picking. Longitudinal conversion junction PCR quantification indicated that converted-ΔSVA_F mESCs remained relatively stable in the population, while converted-ΔSVA_D cells proportion declined moderately compared to XDP cells (**Fig. 1E**). Semi-quantification of PARP1 cleavage, an early apoptosis marker^22–24^, revealed elevated levels in converted-ΔSVA_F cells, and even more so in converted-ΔSVA_D mESCs, suggesting that SVA_F presence is more detrimental (**Fig. 1F**). We reasoned that the failure of isolating converted XDP mESCs is attributed to the additive effect of SVA_F and SVA_D on apoptosis, with SVA_F playing a dominant role.

### SVAs decrease TAF1 expression and impair transcription of TATA-box containing genes

We initially speculated that the human intron-containing hy*TAF1* transcript might be mis-spliced in mESCs in an SVA-dependent manner, leading to non-functional mRNAs. To evaluate this possibility, we amplified the full-length hybrid transcript from mRNA isolated from either transiently converted-XDP mESC pools or converted-ΔSVA_F clones (**Fig. S2A**). Long-read sequencing of these amplicons revealed complete coverage across all exons regardless of species of origin, suggesting no exon skipping during transcription in mESCs (**Fig. S2B**). Furthermore, to rule out potential frameshift mutations missed by pooled sequencing examination, we cloned the amplicons into a TOPO vector and analyzed the sequences of individual bacterial isolates. We found that all captured transcripts were in-frame. Additionally, we detected known (RefSeq) alternative splicing events in exon 5 in the mouse *Taf1* portion, and exons 28’ and 35’ in the human *TAF1* portion (**Fig. S2C**). These data indicate that the hybrid *TAF1* gene can produce canonical transcripts in mESCs.

We next measured hy*TAF1* mRNA levels in converted mESCs using RT-qPCR. Surprisingly, we observed a dramatic reduction in hy*TAF1* mRNA levels in converted mESCs harboring either SVA_F or SVA_D compared with mouse *Taf1* in wild-type mESCs (Fig. 2A). Deleting both SVAs largely restored hy*TAF1* expression, suggesting each SVA independently contributes to the decrease in mRNA abundance (Fig. 2A). Consistent with the mRNA trends, protein level analysis showed markedly reduced TAF1 levels in SVA containing mESCs (Fig. 2B**, Fig. S2D**). Notably, even after deleting both SVAs, hy*TAF1* mRNA and protein levels remained ∼20% below wild type, suggesting that additional *cis-* elements within the human *TAF1* allele may contribute to the reduced expression. To identify such elements, we engineered mESCs in which the human *TAF1* 3′ UTR is swapped back to its mouse counterpart (Fig. 2C), reasoning that 3’ UTRs regulate mRNA stability and translation^26^; or in which all human introns in this region were removed (Fig. 2D). We found that the 3’ UTR swap had no impact on hy*TAF1* mRNA level. In contrast, the intron removal restored expression to near wild-type levels. These experiments confirmed that non-SVA intronic sequences also contribute to a minor extent to the hy*TAF1* expression reduction.

**Figure 2.**
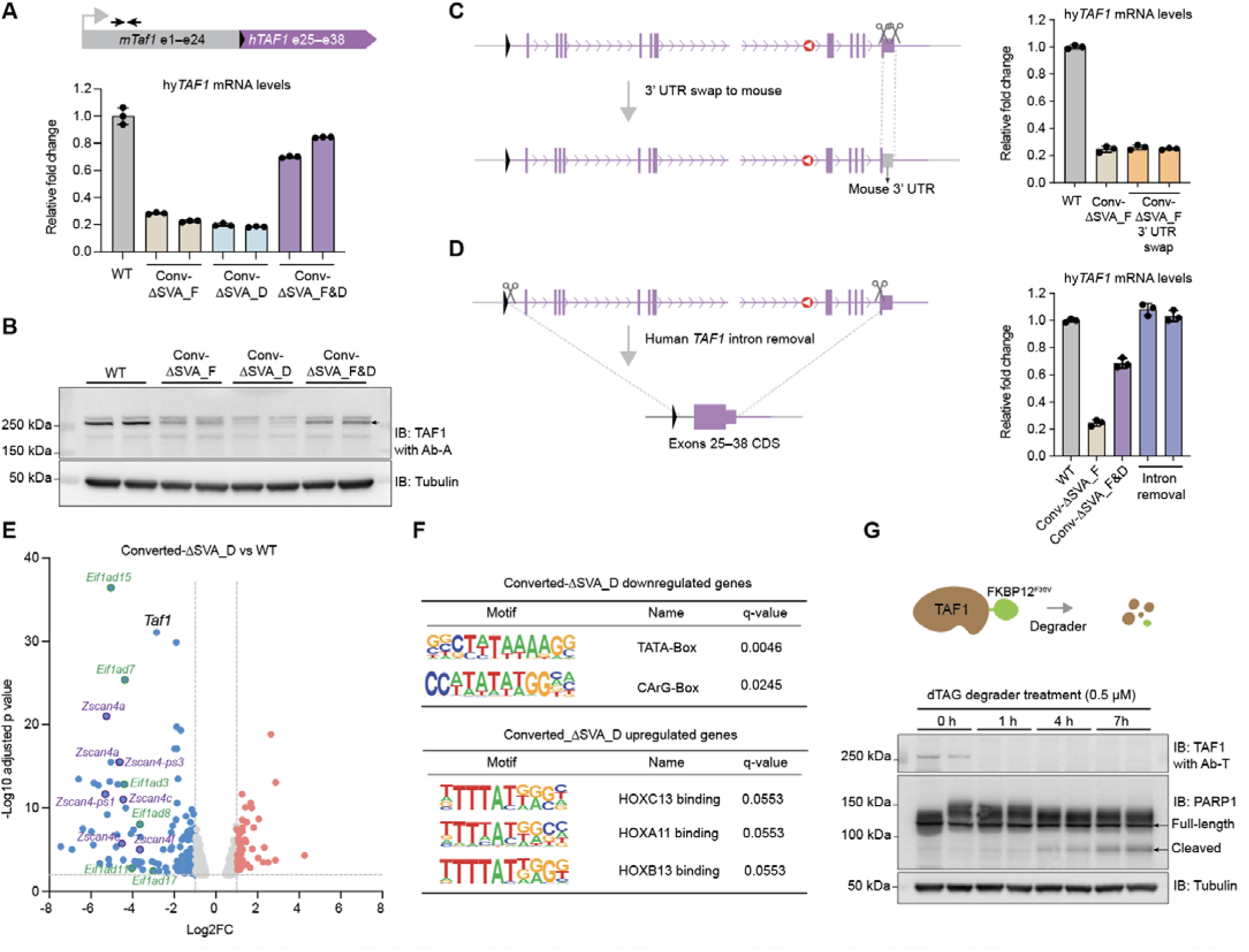
SVAs decrease TAF1 level and impair TATA-box containing gene expression. (A) Quantification of hy*TAF1* mRNA levels in converted cells using primers (oRB617 and oRB618) spanning exons 3–4 (black arrows). Two independent clones were analyzed. Fold change was calculated by normalizing to mouse *Actb* mRNA and to a wild-type clone. Bars represent mean ± SD of three technical replicates. (B) Western blot showing TAF1 expression in converted mESCs. Two independent clones were analyzed. An arrow indicates the TAF1 band. α-Tubulin served as loading control. (**C-D**). Strategies for swapping human *TAF1* 3’ UTR with its mouse counterpart (c) and removing all human introns (d). CRISPR-Cas9 cutting sites are indicated by scissors. A yeast strain harboring the *TAF1*-ΔSVA_F payload was used for *in yeasto* CRISPR editing (see **Methods**) to generate the 3’ UTR swap and the human introns deletion payloads. RT-qPCR analyses of the hy*TAF1* mRNAs are on the right, wild-type, converted-ΔSVA_F and converted-ΔSVA_F&D mESCs were used as controls. Fold change was calculated by normalizing to mouse *Actb* mRNA and to wild-type mESCs. Bars represent mean ± SD of three technical replicates. (E) Volcano plot of differentially expressed genes between converted-ΔSVA_D and wild-type mESCs. Fold change cutoff is 2, adjusted p-value cutoff is 0.01. Red and blue dots represent significantly upregulated and downregulated genes in converted-ΔSVA_D cells, respectively. Highlighted genes in green belong to eukaryotic translation initiation factor 1A family; highlighted genes in purple are involved in telomere maintenance and genomic stability in embryonic stem cells^29^. (F) Motif discovery for differentially expressed genes in converted-ΔSVA_D cells using HOMER^30^. Search regions are from 100 bp upstream to 50 bp downstream of the transcription start site, q-value cutoff is 0.1. (G) Schematic for dTAG-mediated degradation of TAF1 (upper schematic). Western blot analysis of TAF1 and PARP1 after time course degrader treatment. Two independent clones were examined per time point. α-Tubulin serves as loading control.

To assess the functional consequences of hy*TAF1* reduction, we performed RNA-seq and analyzed differential expression between wild-type and converted SVA-containing mESCs. We found that converted-ΔSVA_D cells, which express slightly lower hy*TAF1* level compared to converted-ΔSVA_F cells (Fig. 2A**-B**), exhibit a stronger global transcriptional repression (Fig. 2E**, Fig. S2E**). Gene set enrichment analysis (GSEA) indicated that despite widespread gene repression, only the translation initiation pathway was significantly enriched among downregulated genes in converted-ΔSVA_D mESCs (**Fig. S2F**). In contrast, upregulated transcripts were enriched for gene sets involved in cell differentiation, mitotic progression, and RNA Pol II transcription—potentially indicating compensatory or salvage responses (**Fig. S2F**). Given that TAF1 is a key component of the TFIID complex and is essential for transcription initiation at Pol II promoters, we then analyzed the motif sequences near the transcription start site (TSS) of the differentially expressed genes in converted-ΔSVA_D cells. Notably, TATA box consensus motif was prominent among the downregulated genes (Fig. 2F), implicating compromised TFIID function. Several homeobox (Hox) binding motifs were enriched among the upregulated genes, indicating differentiation of the converted-ΔSVA_D cells (**Fig. S2F**), consistent with previous finding that TFIID complex knockdown compromises mESC pluripotency^27^. Finally, to directly test whether TAF1 depletion is sufficient to induce apoptosis, a phenotype observed in the converted SVA-containing mESCs (Fig. 1F), we fused a dTAG degron^28^ to the C-terminus of the endogenous *Taf1* gene in wild-type mESCs (Fig. 2G top). Treatment with the ligand degrader for 24 hours induced pronounced cell death (**Fig. S2G**). Time-course analysis revealed that TAF1 protein was nearly fully depleted within 1 hour post-treatment. Cleaved PARP1 was elevated at 4 hours and further increased at 7 hours (Fig. 2G bottom), reminiscent of SVAs-containing mESCs (Fig. 1F). In summary, the insertion of either SVA in *TAF1* intron 32 decreases the total hy*TAF1* level and triggers apoptosis, resulting in cell death when both SVA_F and SVA_D are present.

### Chromatin state, transcription of SVAs and their impact on *TAF1* transcription

Transposable elements are rich in regulatory elements^32–34^. To determine the basis of hy*TAF1* reduction caused by SVA insertions, we next investigated the chromatin state of the human *TAF1* region in mESCs. Cleavage under targets and release using nuclease (CUT&RUN)^35,36^ profiling revealed substantial H3K4me3 deposition at both SVA_F and SVA_D (which was accurately detected due to the improved mappability of SVAs in the mouse genome context), indicating open chromatin and active transcription (Fig. 3A). H3K4me3 levels were otherwise indistinguishable between wild-type and humanized mESCs in the regions surrounding the *Taf1* locus (**Fig. S3A**). Stranded RNA-seq revealed that both SVAs are transcribed in the antisense orientation relative to hy*TAF1* and their expression is inversely correlated with hy*TAF1*, particularly of exons 33–38 (Fig. 3B).

**Figure 3.**
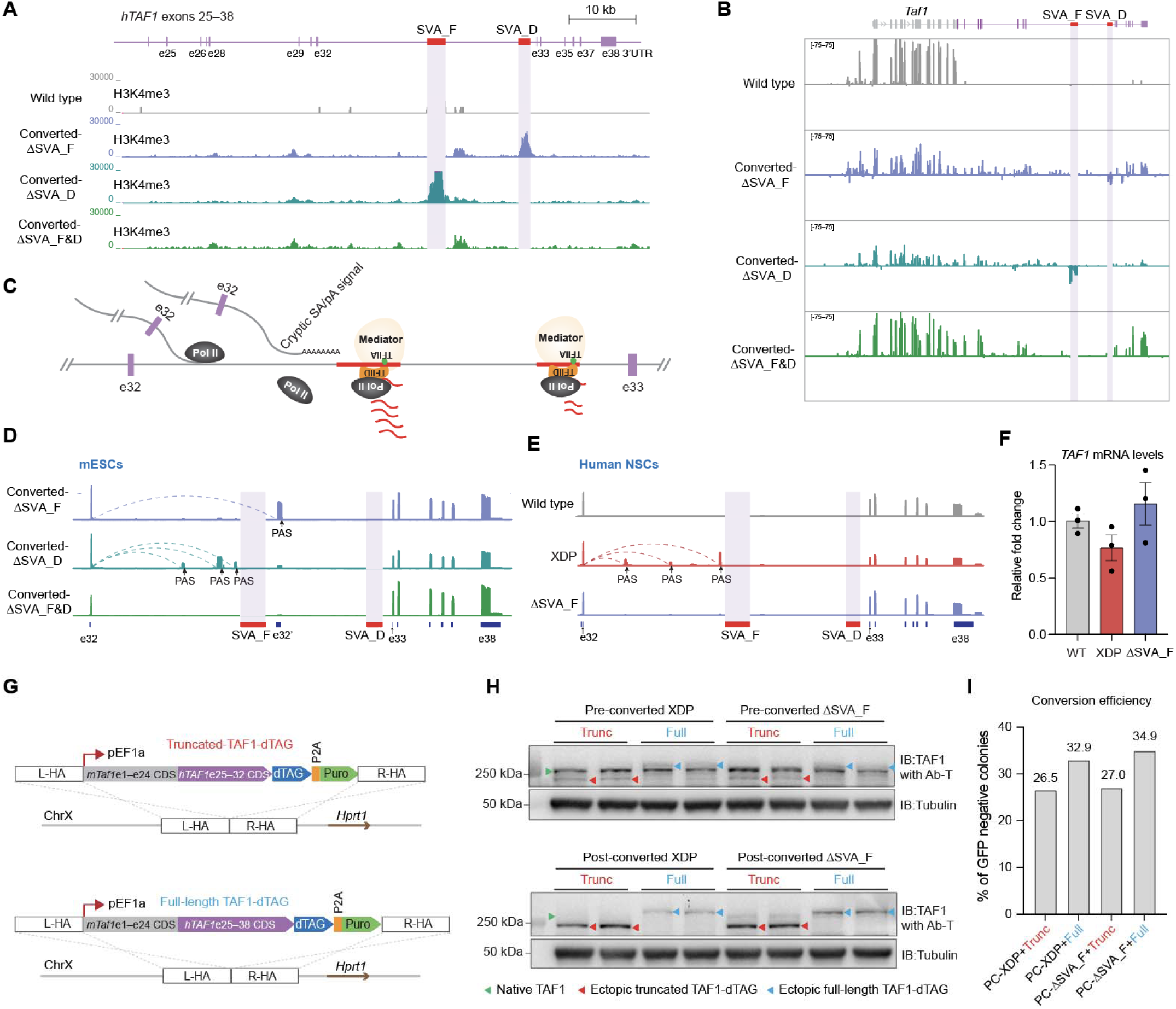
Chromatin state, transcription of SVAs and their impact on *TAF1* transcription. (A) mESCs H3K4me3 CUT&RUN. Sequencing reads were mapped to a custom *TAF1* genomic reference containing human exon 25 through downstream of the 3′ UTR with both SVA_F and SVA_D positioned in intron 32. Transparent purple boxes mark SVA locations. (B) mESCs stranded RNA sequencing. Reads were mapped to a modified mouse reference genome in which *Taf1* exon 25 through the 3′ UTR was replaced with the human *TAF1* counterpart, including both SVA insertions. Transparent purple boxes mark SVA locations. (C) Proposed model: transcription of SVA_F, and to a lesser extent, SVA_D, causes premature termination of *TAF1* transcription at intron 32. Cryptic SA/pA, cryptic splicing acceptor/polyadenylation signal. (D) mESC 3’ RACE using a forward primer targeting human *TAF1* exon 32 and a reverse oligo-dT primer followed by nanopore sequencing (see **Methods** for detail). Dashed arches indicate contiguous reads spanning exon 32 and the cryptic exons. PAS, polyadenylation signal. (E) 3’ RACE sequencing of patient-derived neural stem cells and a ΔSVA_F derivative. Dashed arches indicate contiguous reads spanning exon 32 and the cryptic exons. (F) Quantification of *TAF1* mRNA levels in human neural stem cells by RT-qPCR. Fold change was calculated by normalizing to human *ACTB* mRNA and to a wild-type clone. Bars represent mean ± SEM of three biological replicates. (G) Constructs expressing either truncated (exons 1–32) or full-length hy*TAF1* fused with a dTAG degron targeted to a safe harbor near mouse *Hprt1* on chromosome X. L-HA and R-HA, left and right homology arms, respectively; P2A, porcine teschovirus-1 2A peptide. (H) TAF1 western blot analysis in pre-converted (top) and post-converted (bottom) mESCs expressing either truncated or full-length TAF1-dTAG. α-Tubulin serves as loading control. Two independent mESC clones were analyzed per genotype. (I) Conversion efficiencies for mESCs ectopically expressing either truncated or full-length TAF1-dTAG.

This observation indicates that SVA transcription interferes with transcription elongation of hy*TAF1* mRNA, acting as a transcriptional roadblock. Such interference may cause premature transcription termination and lead to polyadenylation at cryptic polyadenylation sites upstream of the SVAs (Fig. 3C). To test this hypothesis, we performed 3’ rapid amplification of cDNA ends (3’ RACE)^37^ followed by long-read sequencing for converted mESCs (**Fig. S3B**). In converted-ΔSVA_D cells, we detected truncated transcripts containing cryptic exons upstream of SVA_F that terminate at cryptic polyadenylation sites, alongside residual exon 32 readthrough into intron 32 (Fig. 3D**, Fig. S3C**). Similarly, in converted-ΔSVA_F mESCs, a cryptic exon was detected slightly downstream but remained upstream of SVA_D (Fig. 3D, converted-ΔSVA_F). Importantly, exons 33–38 coverage was markedly reduced in SVA-containing cells and was restored only upon double SVA deletion (Fig. 3D, converted-ΔSVA_F&D), supporting the roadblocking model for the SVAs in *TAF1* (Fig. 3D).

To assess the relevance of these findings in a human context, we performed 3’ RACE in XDP patient-derived neural stem cells (NSCs)^15^, along with wild-type and isogenic ΔSVA_F NSCs as controls. We observed similar patterns of premature transcriptional termination in XDP NSCs, which were not observed in the isogenic ΔSVA_F NSCs (Fig. 3E). Furthermore, RT-qPCR confirmed reduced *TAF1* mRNA level in XDP NSCs compared to controls (Fig. 3F).

Given that TAF1 functions as part of the TFIID complex by interacting with other TAF proteins^16^, we asked whether a truncated TAF1 generated by premature termination could exert dominant-negative effects. To address this, we integrated either full-length or truncated (exons 1–32) dTAG-tagged hy*TAF1* coding sequence into an X chromosome safe harbor locus of pre-converted-XDP and pre-converted-ΔSVA_F mESCs (Fig. 3G). We found that the truncated hyTAF1 and the native mouse TAF1 coexist in the pre-converted mESCs (Fig. 3H, top panel, red and green arrow, respectively) and those mESCs grew normally (**Fig. S3D-E**), arguing against a potential dominant-negative role of the truncated TAF1. Interestingly, upon Cre delivery, we found even the XDP mESCs with ectopic truncated hyTAF1 could convert efficiently (Fig. 3I**, Fig. S3M**). When examining the TAF1 protein in post-converted clones, native TAF1 level was strongly diminished relative to wild type, particularly in XDP mESCs, whereas truncated hyTAF1 expression persisted or even increased (Fig. 3H, bottom panel). These results indicate that the truncated hyTAF1 is not only non-deleterious to cell viability but may in fact provide partial compensation in the context of native TAF1 depletion.

Structural modeling of TAF1 revealed that exons 33–38 largely encode disordered regions, with only a short α-helical domain encoding the second half of the second bromodomain, raising the possibility that these C-terminal exons might be functionally dispensable (**Fig. S3F**). To directly test this hypothesis, we first deleted the region spanning intron 32 (including both SVAs) through the stop codon (leaving the 3’ UTR intact) in pre-converted XDP mESCs, followed by Cre-mediated conversion (**Fig. S3G**). Converted clones only expressing hy*TAF1* exons 1–32 were readily detected (**Fig. S3H-I**). However, these clones expressed reduced levels of hy*TAF1* e1–32 (**Fig. S3J-K**) and proliferated more slowly compared to wild-type mESCs (**Fig. S3L**). These findings suggest that the truncated hy*TAF1* transcript may be unstable and subject to degradation, and that residual truncated hy*TAF1* expression can partially support cell viability.

Taken together, SVA insertions in *TAF1* intron 32 establish a chromatin and transcriptional environment that promotes active antisense transcription and obstructs productive elongation of the hy*TAF1* transcript. Alternatively, the propensity for G-quadruplex formation in the SVA might impede hy*TAF1* transcription through the SVAs^38,39^. Interference with hy*TAF1* transcription results in premature termination, and most of the prematurely terminated hy*TAF1* transcripts are presumably subjected to degradation, while residual prematurely terminated hy*TAF1* may produce a truncated yet partially functional TAF1 protein.

### A conditional XDP mouse model recapitulates disease phenotypes

Due to the detrimental effect of the SVAs on *TAF1* transcription, we were unable to isolate viable converted XDP mESCs to directly establish a disease mouse model. Instead, we generated conditional XDP mice by injecting the pre-converted XDP mESCs into wild-type blastocysts and obtained chimeric male mice first. Chimeric male founders were bred to wild-type females to obtain heterozygous pre-converted XDP females, which were then crossed with homozygous nestin-Cre (*Nes*^Cre^) males (Fig. 4A). This strategy enables Cre expression in neuronal and glial precursors^40^, thereby conditionally inducing hy*TAF1* expression in the central and peripheral nervous system of the progeny. Among the 144 male progeny genotyped, 70 carried the *Nes*^Cre^hy*TAF1*^XDP^ allele while 74 did not, consistent with Mendelian segregation (χ^2^ = 0.111, p = 0.74, chi-square test), indicating that the *Nes*^Cre^hy*TAF1*^XDP^ allele does not cause lethality at two weeks of age (genotyping time point) in males^41^. However, all *Nes*^Cre^hy*TAF1*^XDP^ males died by two months of age, whereas littermate *Nes*^Cre^ control males exhibited normal survival (Fig. 4B) and no gross phenotype. Although the affected *Nes*^Cre^hy*TAF1*^XDP^ males had reduced body size compared to wild type control, no obvious skeleton abnormality was detected in examined mice by X-ray absorptiometry (Fig. 4C). Heterozygous females, which undergo random X-chromosome inactivation, were largely unaffected for survival (Fig. 4B). In contrast, homozygous XDP females exhibited lethality and reduced body size similar to males (Fig. 4B**, Fig. S4A**), indicating that XDP allele dosage underlies the phenotype, and that the mouse model recapitulates human genetics^42^.

**Figure 4.**
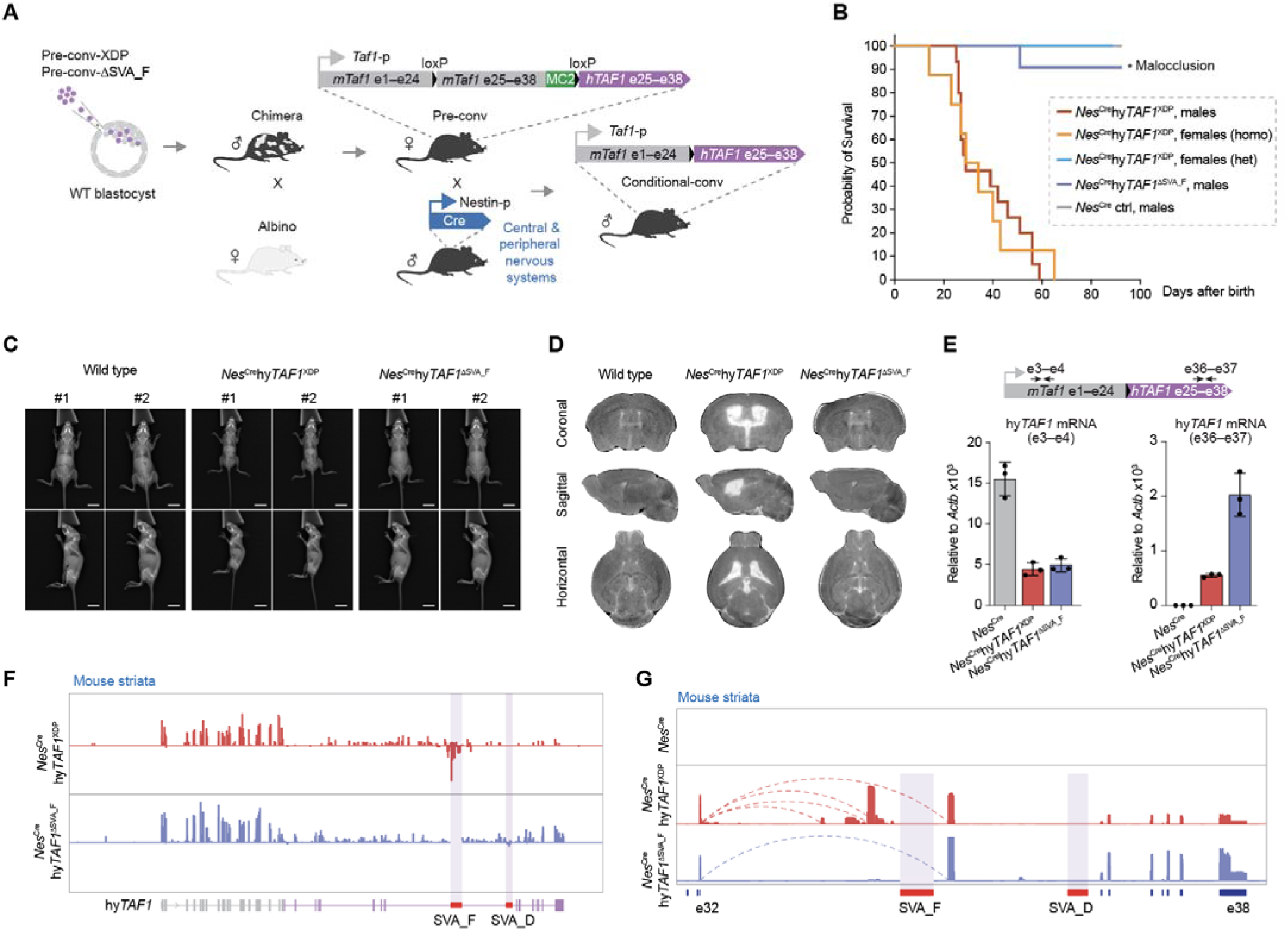
A conditional XDP mouse model recapitulates disease phenotypes. (A) Breeding scheme for generating neuronally converted hy*TAF1* male mice. Founder male mice carrying the pre-converted (pre-conv) XDP allele were derived by chimeric blastocyst injection and crossed with a wild-type albino C57BL/6N female to obtain heterozygous females, which were subsequently bred with C57BL/6J nestin-Cre males to drive neural-specific conversion in male progeny. (B) Kaplan–Meier survival curves of *Nes*^Cre^hy*TAF1*^XDP^ males, *Nes*^Cre^hy*TAF1*^ΔSVA_F^ males, heterozygous *Nes*^Cre^hy*TAF1*^XDP^ females and homozygous *Nes*^Cre^hy*TAF1*^XDP^ females. *Nes*^Cre^ littermates were included as controls. Asterisk marks a *Nes*^Cre^hy*TAF1*^ΔSVA_F^ male that died from malocclusion, an unrelated phenotype occasionally observed in inbred strains. (C) Representative images for dual-energy X-ray absorptiometry (DEXA) scan of wild type, *Nes*^Cre^hy*TAF1*^XDP^ and *Nes*^Cre^hy*TAF1*^ΔSVA_F^ male mice. Two mice per group were shown. All mice were scanned on postnatal day 22 with dorsal and lateral views. Scale bar is 1 cm. (D) Representative MRI images of matched brain sections from postnatal day 22 wild type, *Nes*^Cre^hy*TAF1*^XDP^ and *Nes*^Cre^hy*TAF1*^ΔSVA_F^ male mice, highlighting pronounced striatal atrophy in *Nes*^Cre^hy*TAF1*^XDP^ brains. (E) Quantification of hy*TAF1* mRNA in conditionally converted 3-week-old mouse striata by RT-qPCR. Primer pairs spanning both mouse and human exon junctions were used (black arrows). Bars represent mean ± SD of three biological replicates. (F) Stranded RNA sequencing of striatal samples from conditionally converted mice. Reads were mapped to a custom mouse reference genome in which *Taf1* exon 25 through the 3′ UTR was replaced with the human *TAF1* region carrying both SVAs in intron 32. Transparent purple boxes indicate SVA insertion sites. (G) 3′ RACE sequencing of striatal RNA from conditionally converted animals. Prematurely terminated hy*TAF1* transcripts are observed in *Nes*^Cre^hy*TAF1*^XDP^ but absent in *Nes*^Cre^hy*TAF1*^ΔSVA_F^ striata. Dashed arches indicate contiguous reads spanning exon 32 and the cryptic exons.

To determine whether SVA_F is the major driver of this severe pathology, we generated conditional ΔSVA_F mice using the pre-converted-ΔSVA_F mESCs (Fig. 1C) and followed the same breeding scheme (Fig. 4A). Remarkably, conditionally converted-ΔSVA_F male progeny all survived beyond three months (Fig. 4B) and continue to live without detectable abnormalities, except slightly lower body weight comparing to wild type (Fig. 4C**, Fig. S4B**).

MRI scanning of three-week-old *Nes*^Cre^hy*TAF1*^XDP^ male brains revealed pronounced striatal atrophy coupled with enlarged lateral ventricles—a phenomenon absent in the *Nes*^Cre^hy*TAF1*^ΔSVA_F^ male mice (Fig. 4D**, Fig. S4C**). Notably, nestin-Cre-mediated conversion efficiencies in both *Nes*^Cre^hy*TAF1*^XDP^ and *Nes*^Cre^hy*TAF1*^ΔSVA_F^ striata exceeded 80% (**Fig. S4D**), suggesting that the severe phenotype in *Nes*^Cre^hy*TAF1*^XDP^ mice is driven by SVA_F, rather than by differences in conversion efficiencies.

Given the role of repeat expansion in many other neurodegenerative diseases^43^, and prior observations linking XDP SVA_F hexamer repeat length with age of onset in XDP patients^38^, we wondered whether repeat expansion occurred in the XDP mouse brain. Amplicons of the SVA_F hexamer region from striatal and cortical genomic DNAs showed no evidence of somatic expansion by gel electrophoresis (**Fig. S4E**). We also cloned the striatum- and cortex-derived amplicons into a TOPO vector and sequenced five bacterial isolates each. All isolates had consistent 34 contiguous hexamer repeats, exactly matching the parental XDP BAC used for *TAF1* humanization (**Fig. S4E**). Thus, bulk hexamer repeats are stable at the examined age, though this does not rule out some level of expansion in a subset of cells which might be short lived^43^.

To evaluate the consequences of the SVA_F insertion *in vivo*, we quantified the level of bulk striatal *Taf1* mRNA, which is mostly transcribed by converted hy*TAF1*, as well as from a small proportion of non-converted cells. Both *Nes*^Cre^hy*TAF1*^XDP^ and *Nes*^Cre^hy*TAF1*^ΔSVA_F^ mice exhibited reduced total *Taf1* levels compared to the wild-type controls (Fig. 4E). However, when probing the human *TAF1* exon 36–37 junction, indicative of successful transcriptional pass through of intron 32, *Nes*^Cre^hy*TAF1*^ΔSVA_F^ mice had significantly higher levels than *Nes*^Cre^hy*TAF1*^XDP^ mice, consistent with transcriptional interference by SVA_F (Fig. 4E). Stranded RNA-seq of striatal RNA confirmed active antisense transcription from SVA_F and decreased coverage of downstream *TAF1* exons in *Nes*^Cre^hy*TAF1*^XDP^ but not in *Nes*^Cre^hy*TAF1*^ΔSVA_F^ striatum (Fig. 4F**, Fig. S4F**). Similarly, 3’ RACE of striatal RNA showed prematurely terminated transcripts upstream of SVA_F specifically in *Nes*^Cre^hy*TAF1*^XDP^ striata (Fig. 4G).

To assess whether this phenomenon is conserved in humans, we performed 3’ RACE on caudate nucleus RNA from three postmortem XDP patients and one unaffected control^44^. In all XDP samples, prematurely terminated *TAF1* transcripts were detected near cryptic polyadenylation sites within intron 32 (**Fig. S4G**), consistent with the pattern seen in *Nes*^Cre^hy*TAF1*^XDP^ mice (Fig. 4G). Collectively, these findings demonstrate that SVA_F is sufficient to block transcriptional readthrough of intron 32 and drive striatal atrophy in a humanized XDP mouse model. The conditional XDP mouse model recapitulates key molecular and pathological features observed in XDP patients as is further described in detail in the accompanying paper^41^.

### Transcriptional activation of *Taf1* rescues XDP lethality

Next, we sought to explore potential therapeutic strategies for XDP using our hy*TAF1* model. Notably, deleting SVA_F enabled efficient mESC conversion (Fig. 1D), although the converted-ΔSVA_F mESCs express only ∼25% of the wild-type *Taf1* levels (Fig. 2A). Yet this reduced expression is sufficient to support nearly normal cell proliferation rate (**Fig. S5A**). *In vivo*, although striatal hy*TAF1* expression was significantly reduced (Fig. 4E), *Nes*^Cre^hy*TAF1*^ΔSVA_F^ mice showed much improved survival, increased body size and absence of striatal atrophy (Fig. 4B**-D**) compared to *Nes*^Cre^hy*TAF1*^XDP^ mice, suggesting tolerance to substantially reduced hy*TAF1* level. Cell death occurs only when full-length hy*TAF1* levels drop below a critical threshold (due to the additive effect of both SVAs). These observations are consistent with transcriptomic analyses of postmortem human caudate nuclei, which show *TAF1* expression declines steadily after age 15, with a more pronounced drop around age 40^45,46^, which correlates with the average age of XDP onset. Based on this threshold model, we hypothesize that XDP symptoms could be delayed or ameliorated by moderately boosting *TAF1* expression using targeted gene activation strategies. As a proof of concept, we employed a CRISPR activation (CRISPRa) system in converted-ΔSVA_D mESCs, which retain the pathogenic SVA_F element and express only ∼20% of wild-type *Taf1* levels (Fig. 2A). Four gRNAs, with binding sites ranging from 23 to 311 bp upstream of the *Taf1* transcription start site (TSS), were cloned under a U6 promoter into a VPR-dCas9 expression vector^47^ (Fig. 5A). The VPR-dCas9-gRNA cassette was stably integrated into the converted-ΔSVA_D mESCs via a piggyBAC transposon system and dCas9 expression was confirmed by western blot analysis (Fig. 5B). Quantification of the endogenous *Taf1* mRNA showed restoration of expression: VPR-dCas9-gRNA (−23) and -gRNA (−311) increased hy*TAF1* to ∼40% of wild-type levels, while VPR-dCas9-gRNA (−246) restored expression to ∼50% (Fig. 5C left). Importantly, quantification of the human *TAF1* portion confirmed upregulation but to a lesser extent, suggesting persistent transcriptional barrier caused by SVA_F (Fig. 5C right). Nevertheless, given that even the truncated hy*TAF1* is partially functional (**Fig. S3F-L**), we therefore hypothesized that CRISPRa could enable conversion of XDP mESCs. To test this, the VPR-dCas9-gRNA (−246) expression plasmid was stably integrated into pre-converted XDP mESCs via piggyBAC transposon system, followed by Cre-mediated conversion. Longitudinal conversion junction PCR quantification showed that VPR-dCas9-gRNA (−246) slowed down the depletion of converted cells in the population (Fig. 5D). Approximately 17.9% of the VPR-dCas9-gRNA (−246) colonies and 6.6% of the VPR-dCas9-no gRNA colonies were GFP negative (Fig. 5E**, Fig. S5B**). Importantly, the VPR-dCas9-no gRNA converted cells formed significantly smaller colonies (Fig. 5F) and failed to recover after expansion, whereas VPR-dCas9-gRNA (−246) converted-XDP colonies survived after expansion. Genotyping PCR confirmed successful conversion and the presence of both SVAs in all the isolated clones (Fig. 5G). Notably, VPR-dCas9-gRNA (−246) rescued the hy*TAF1* mRNA levels to variying degrees in these clones, with the highest reaching ∼70% of the wild type level (Fig. 5H). Given that piggyBAC transposon-mediated integration can result in expression difference due to position effect, we quantified *dCas9* mRNA levels and observed a positive correlation with hy*TAF1* mRNA abundance (Fig. 5I). Together, these data suggest boosting *Taf1* transcription could ameliorate XDP phenotypes, providing a feasible solution for treating XDP.

**Figure 5.**
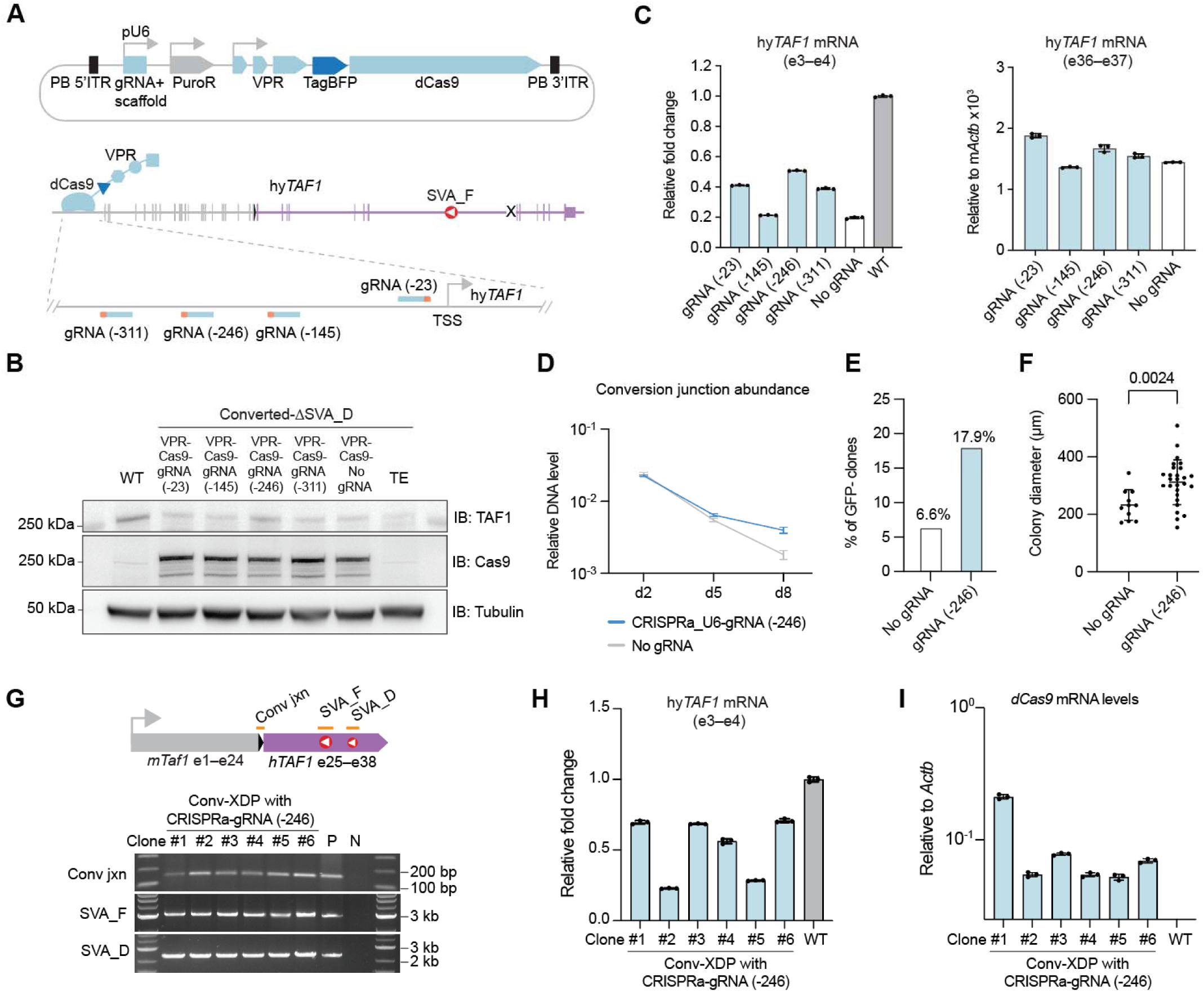
Transcriptional activation of hy*TAF1* rescues XDP lethality. (A) Strategy for CRISPRa-mediated boosting of hy*TAF1* transcription. Upper panel shows the components of the CRISPRa cassette flanked by the inverted terminal repeats of the piggyBAC transposon. Bottom panel shows the positions of the gRNAs relative to the TSS of the hy*TAF1* gene. (B) Western blot analysis of dCas9 in converted-ΔSVA_D mESCs. mESCs were first co-transfected with piggyBAC transposase expression plasmid (System Biosciences) and VPR-dCas9 cargo plasmid, followed by puromycin selection until all the cells gained resistance. TE, converted-ΔSVA_D cells transfected with TE buffer. (C) RT-qPCR quantification of hy*TAF1* mRNA levels using primers spanning exon 3–4 (left panel). Fold change was calculated by normalizing to mouse *Actb* and to wild-type mESCs. mESCs with VPR-dCas9 only (no gRNA) serve as baseline control. mRNA quantification using primers spanning exon 36–37 (right panel) indicates the readthrough transcript levels, with relative levels calculated by normalizing to mouse *Actb*. Bars represent mean ± SD of three technical replicates. (D) Longitudinal quantification of conversion junction abundance. A genomic mouse *Actb* primer pair was used as an internal reference. (E) Percentages of GFP negative colonies after Cre-mediated conversion. VPR-dCas9-no gRNA served as control. (F) Diameter of GFP negative colonies. Two-tailed Mann-Whitney *U* test. (G) Genotyping PCR of single-colony isolated converted-XDP mESCs. Orange bars mark the locations of conversion junction, full-length SVA_F and SVA_D. P, positive control from transiently converted XDP mESCs pool; N, negative control from wild type mESCs. (H) hy*TAF1* mRNA levels in converted-XDP clones. Bars represent mean ± SD of three technical replicates. (I) *dCas9* mRNA levels in converted-XDP clones. Bars represent mean ± SD of three technical replicates.

## Discussion

SVA retrotransposons represent a primate-specific family of mobile elements that are active in humans and have emerged as an important contributor to a variety of human genetic diseases, including the founder effect disease Fukuyama muscular dystrophy, which is characterized by aberrant splicing^11^. Indeed, additional genetic disorders associated with SVA insertions have been characterized and are associated with a panoply of altered pre-mRNA splicing patterns in the affected genes^48–50^. Mis-splicing has also been implicated in a founder *Alu* retrotransposon insertion that led to a trait shared among all great apes—loss of the tail^51^. Thus, founder effect mutations caused by retrotransposons can have lasting effects on populations, as appears to be the case in XDP, which originated in the Philippines^52^.

At present, there is no effective cure for XDP; although many patients are treated with medications for general dystonia, such as anticholinergic agents, muscle relaxants and other benzodiazepines^52,53^. These treatments only offer temporary or partial relief and inconsistent benefits. Discovering effective therapies is thus an urgent need. In this study, we report the creation of a humanized mouse model for XDP by carefully replacing a 3’ segment of the mouse *Taf1* gene with an XDP patient-derived human *TAF1* gene, including the disease-causing SVA_F insertion, without disturbing the 5’ regulatory elements and 5’ half of the gene body. *In vitro*, we show that intronic SVA elements reduce gene expression by forming transcriptional roadblocks, leading to prematurely polyadenylated transcripts and insufficient TAF1 protein. This mechanism appears conserved in cells of both mouse (mESCs) and human (hNSCs). Notably, minor H3K4me3 enrichment and transcription were also detected in human intron 28 and the i32′ region of *TAF1* (Fig. 3A**-B**), which may explain the result that hy*TAF1* mRNAs remained ∼20% below wild type after deleting both SVAs in intron 32 (Fig. 2A**, 2D**)*. In vivo*, Cre-mediated formation of hy*TAF1* in the developing mouse nervous system leads to rapid lethality accompanied by striatal atrophy, which is the hallmark phenotype of XDP. In contrast, the isogenic ΔSVA_F model was phenotypically near normal, directly implicating the SVA_F insertion as the molecular driver^15^. Notably, XDP mice develop symptoms up to 300 times faster than human patients, which is likely due to the absence of human-specific ZNF91^8,19^, which binds SVAs and mediates their transcriptional repression. Similar to the results seen in mESCs and hNSCs, SVAs are actively transcribed in the mouse brain, blocking hy*TAF1* transcription. Besides the transcriptional activity of SVAs, DNA G-quadruplexes within the VNTR region^38^ may act as another barrier to RNA Polymerase II-mediated elongation^39^. These findings highlight the value of preserving human intronic architecture for modeling diseases in which pathogenesis arises from noncoding genomic regions.

Beyond modeling, our system provides a platform for therapeutic development. We showed that transcriptional activation of the hy*TAF1* allele using CRISPRa partially rescues gene expression and improves cellular viability *in vitro*. This proof-of-concept experiment establishes a rationale for developing smaller targeted gene activators, such as artificial zinc finger proteins^54^ that bind the promoter of hy*TAF1*, and can be packaged into an adeno-associated virus (AAV) capsid for efficient delivery into the mouse brain. Although the human *TAF1* promoter sequence is different from that of mouse, our mouse model now enables *in vivo* testing of AAV-based strategies to boost hy*TAF1* expression, evaluating delivery route, safety and efficacy. Given the well-established use of AAVs in the clinic^55–57^, this model provides a tangible framework for preclinical studies.

One of the innovations of this work is the application of large-scale mouse genome rewriting to model a complex human allele with non-coding pathogenic elements—the SVA retrotransposons. The conditional conversion design circumvents the challenge where directly inserting the SVAs is lethal to cells. More broadly, the mouse genome rewriting approach described here exemplifies the study of human-specific noncoding variants underlying rare genetic disorders. As genome-wide association studies (GWAS) and long-read sequencing continue to reveal roles of transposable elements in diseases, scalable strategies for incorporating these elements into model organisms will become increasingly necessary.

Together, our work establishes a generalizable strategy for building accurate mouse models of human rare disease and demonstrates its utility in elucidating pathogenic mechanisms and enabling therapeutic discovery. The XDP humanized mouse model showcases the power of genomic tailoring not only to recapitulate disease biology in mice, but also to serve as a therapeutic testing ground for translational advances beneficial to humans.

## Methods

### Strains and plasmids

Yeast BY4741 strain was used for all yeast assembly and editing. A mouse embryonic stem cell line derived from inbred C57BL/6J mice was from NYU Langone Health Rodent Genetic Engineering Laboratory. Albino C57BL/6N female mice were purchased from Charles River Laboratories (strain 562). Nestin-Cre C57BL/6J mice were purchased from the Jackson Laboratory (strain 003771). All animal experimental procedures were approved by the Institutional Animal Care and Use Committee (IACUC) at NYU Langone Health. Human caudate nuclei, neural stem cells and XDP patient-derived BACs (XDP BAC AEX253) spanning the *TAF1* gene and its flanking sequences were obtained from the Collaborative Center for XDP (CCXDP). Mouse BAC RP23-334A11 and human BAC RP11-554L2 were purchased from the BACPAC Resources Center. Marker cassette 1 donor plasmid was cloned by assembling two homology arms and marker cassette 1^18^ into a linearized pUC19 vector using the NEBuilder HiFi DNA Assembly Master Mix (NEB, E2621X). Yeast Cas9 and guide RNA expression plasmids are from Zhao *et al.*^25^.

### Payload DNA assembly in yeast

XDP patient-derived bacterial artificial chromosome (BAC) AEX253 was extracted using a BAC DNA purification kit (Takara, 740436.25) by following the manufacturer’s instructions and digested using the *in vitro* CRISPR system (Cas9 nuclease, NEB, M0386T; two custom crRNA:tracrRNA duplexes were ordered from IDT Inc.). The sequences and coordinates for the two custom crRNAs (*hTAF1*_1 and *hTAF1*_2) are listed in Supplementary Table 2. Digested BAC DNA was co-transformed with a linear acceptor vector into the BY4741 yeast cells using a previously described protocol^58^. The linear acceptor vector contained two 200-bp homology arms corresponding to the termini of the digested BAC, directing homologous recombination in yeast. Transformants were selected on Synthetic Complete (SC)–Ura plates and PCR-screened for presence of two novel assembly junctions. Newly assembled payload DNA was extracted from yeast using the yeast miniprep kit (Zymo Research, D2001), and was subsequently electroporated into the EPI300 *E. coli* cells (Lucigen, EC300150) for plasmid amplification under induction condition (Lucigen, CCIS125). Payload DNA was extracted from *E. coli* using the BAC DNA purification kit and sequenced on an Illumina 550 sequencer for verification. Sequencing data was inspected as described previously^21^.

### Payload DNA editing in yeast

Yeast cells carrying an episomal payload were inoculated in 5 ml SC–Ura medium overnight. In the following day, yeast culture was diluted in fresh SC–Ura medium with starting A_600_=0.15. Yeast cells were harvested after 4 hours of growth at 30 °C for competent cells preparation. *Sp*Cas9 was driven by the *Sc*PGK1 promoter, sgRNAs were driven by a tRNA gene promoter and expressed from a 2-micron plasmid. sgRNA sequences are in **Supplementary Table 2**. Repair donor DNAs were either synthesized by IDT or generated by fusion PCR. 100 ng *Sc*PGK1p-*Sp*Cas9-sgRNA and 50 ng donor DNA were co-transformed into the yeast competent cells harboring the payload. Transformants were plated on SC–Ura–His medium for two days of incubation at 30 °C. Successfully edited yeast clones were confirmed by Sanger sequencing of the edited region(s). All final constructs were also verified using Illumina sequencing as described above.

### mESCs engineering

Three types of media were used: 1. mES medium for culturing mESCs on feeder cells contained KnockOut DMEM (Fisher Scientific, 10829018), 15% Fetal Bovine Serum (GE Healthcare Life Sciences, SH30070.03), 0.1 mM MEM Non-Essential Amino Acids (Fisher Scientific, 11140050), 1% Penicillin-Streptomycin-Glutamine (Invitrogen, 10378-016), 1000 U/ml LIF (EMD Millipore, ESG1107), 0.2 mM 2-Mercaptoethanol (7 µl pure solution into 500 ml). 2. ESM+2i medium for culturing mESCs on gelatin contained mES medium supplemented with 3 µM of CHIR 99021 (Tocris, 4423) and 1 µM MEK inhibitor (STEMCELL Technologies, 72184). 3. MEF medium for seeding mouse embryonic fibroblasts (MEFs) feeder cells contained DMEM (Life Technologies, 11965-118), 10% Fetal Bovine Serum (Gemini Bio-Products), 0.1 mM MEM Non-Essential Amino Acids, 1% Penicillin-Streptomycin-Glutamine. Mitotically inactivated MEF feeder cells (Cell Biolabs, CBA-310, mitotically inactivated by mitomycin C treatment following a previously described protocol^59^) were seeded at 5–7.5 × 10^4^ cells/cm^2^ on 0.1% gelatin-coated plates one day prior to mESCs seeding. MEF medium was replaced with mES medium at least 2 hours before thawing and seeding ∼1 million mESCs per 6 cm plate. mESCs were cultured in mES medium on feeders with daily medium changes and passaged at a 1:3 ratio upon reaching 70–80% confluency.

For nucleofection, 2–3 million mESCs were electroporated (Mirus Bio MIR50118; Lonza Nucleofector 2b, program A-023) with 5 µg payload DNA and 2 µg Cas9-gRNA plasmids for *hTAF1* payload delivery, or 0.5 nmol single-stranded oligodeoxynucleotide and 2 µg Cas9-gRNA plasmid for loxP insertion into *Taf1* intron 24. Electroporated mESCs were seeded with the appropriate drug-resistant feeder cells (EMD Millipore, PMEF-NL or Applied StemCell, ASF-1001). Medium was changed 12–16 hours post-nucleofection, and antibiotic selection was applied 24–36 hours post-nucleofection. Antibiotic concentrations used included: puromycin, 0.8 µg/ml; geneticin, 150 µg/ml. Colonies were monitored daily until reaching appropriate sizes. Colonies were picked into round-bottom low retention 96-well plates and trypsinized with 0.25% Trypsin-EDTA (Gibco, 25200056) at 37 °C for 7 minutes and neutralized with 100 µl ES medium. Cells were singularized by manually pipetting 10–20 times and split between gelatin-coated plate and feeder-coated 96-well plates. Media were refreshed daily. Once over 50% confluency was reached for the gelatin-coated plate, ∼10% of cells were passaged for expansion and the remainder used for crude genomic DNA extraction and PCR genotyping. Once correctly edited clones were identified, the corresponding cells growing on feeders were stocked in freezing medium (90% FBS, 10% DMSO) in liquid nitrogen. Unless otherwise specified, mESCs were cultured at 37 °C in a humidified incubator with CO_2_ set to 5%.

### Human neural stem cells differentiation

Collection and reprogramming of XDP patient and control samples into induced pluripotent stem cells (iPSCs), as well as CRISPR-Cas9 targeting for SVA_F removal were described previously^15,60^. Neural stem cells (NSCs) derived from XDP, control and ΔSVA_F iPSCs were generated using a PSC Neural Induction medium (Gibco, A1647801), described in detail in Aneichyk *et al*^15^. Briefly, iPSCs were plated as single cells on Geltrex (Thermo Fisher Scientific, 12760013) coated dishes and cultured in Neural Induction Medium for 7 days. Cells were collected and plated onto Geltrex coated dishes in Neural Expansion Medium (50% Neural Induction Medium, 50% Advanced DMEM/F-12 (Gibco, 12634028)) with rock inhibitor (Y-27632; 5 μM). Fresh Expansion medium was replaced after 24 hours to remove rock inhibitor, and cells were grown to full confluency and stored as NSC passage 1. NSCs were frozen at passages 3 and 4 at −80 °C for further use.

### hy*TAF1* mESC variants engineering

hy*TAF1* variants including ΔSVA_F, ΔSVA_D, 3′ UTR replacement, and human *TAF1* intron removal were first constructed in yeast by editing the initial *hTAF1*-XDP payload for generating the corresponding variant payloads. All variant payloads were confirmed by Illumina sequencing prior to introduction into mESCs. Resulting mESC lines were validated by targeted sequencing (see below) of the integration site and introduced DNA plasmids.

### Genomic DNA extraction

Crude genomic DNA from mESCs was extracted by lysing mESC pellets in 30 μl of lysis buffer (0.3 mg/ml proteinase K) at 50 °C for 1 hour, followed by 98 °C 10[minutes. Genomic DNAs used for targeted sequencing library preparation, conversion junction quantification, DNA copy number quantification, full-length SVA_F amplification, and hexamer repeat length evaluation experiments were extracted from either mESCs or mouse brain tissues using a QIAamp DNA Mini Kit (QIAGEN, 51306).

### hy*TAF1* Conversion and quantification

Approximately 1 µg pCAG-iCre plasmid (Addgene, 89573) was nucleofected into 2 million pre-converted mESCs using the protocol mentioned above. Nucleofected cells were plated in a 6-well plate. ESM+2i medium was refreshed daily. Cells were passaged in a 1:10 ratio every 3 days, 90% of the cells were harvested for genomic DNA extraction at each time point. Genomic DNA samples were quantified using a Qubit dsDNA board range assay kit (Thermo Scientific, Q33266). Approximately 100 ng genomic DNA was used in a 10 µl SYBR Green (Roche, 04887352001) quantitative PCR (qPCR) reactions (see below). Primers targeting the conversion junction (oWZ2213 and oWZ2214) and primers targeting the endogenous *Actb* intronic region (oWZ1683 and oWZ1684) were used. Conversion was estimated by normalizing the conversion junction DNA abundance to *Actb*.

### mESCs imaging

For conversion experiments, following the Cre plasmid nucleofection into pre-converted mESCs, approximately 5,000 cells were seeded in a 6-well plate and continuously monitored using an Incucyte S3 instrument for 8 days, medium was swapped daily. Images were taken every 6 hours. At the final time point, single colonies were scanned using an ALS CellCelector instrument or an EVOS 5000 (Invitrogen, AMF5000) instrument under both bright field and GFP channels.

### Mouse genotyping

Genomic DNA was extracted from mouse tail or ear biopsies by lysis in 30 µl DirectPCR Lysis Reagent (Viagen Biotech, 102-T) supplemented with 30 µg proteinase K. Samples were incubated at 55[°C for 3 hours, followed by heat inactivation at 95[°C for 10[minutes. After a brief centrifugation, 1 µl of the supernatant was used as template in a 10 µl PCR reaction containing 5 µl GoTaq Green Master Mix (Promega, M7123) and 2 µM each of forward and reverse primers (**Supplementary Table 1**). The PCR program was as follows: 95[°C for 5[minutes; 30 cycles of 95[°C for 30 s, 58[°C for 30 s, and 72[°C for 30 s; 72[°C for 5[minutes.

### Whole-body composition analysis

Whole-body composition was assessed using a dual-energy X-ray absorptiometry (DEXA) Insight scanner (OsteoSys, Scintica), harboring a single X-ray source adjustable between 40 and 80 keV. Three-weeks-old mice were anesthetized in an isoflurane induction chamber for approximately 3 minutes, with anesthesia confirmed via toe pinch. Animals were then positioned into the scanner’s heated animal stage to maintain body temperature and minimize motion artifacts. Isoflurane (1.5–2.5%) in oxygen (1 L/min) was continuously administered via an integrated nose cone throughout the imaging session. Each mouse underwent two scans—dorsal and lateral views—to obtain measurements including total mouse body weight, fat and lean mass, bone mineral density (BMD), bone mineral content (BMC), bone area, tissue area were recorded and exported for analysis.

### MRI preparation and mouse brain imaging

At experimental endpoints, mice were deeply anesthetized (ketamine 100 mg/kg, xylazine 10 mg/kg) and transcardially perfused first with PBS followed by 4% paraformaldehyde (PFA) using a low flow rate minipump. The decapitated heads were then stored in 4% PFA at 4[°C for at least 48 hours. High-throughput *ex vivo* MRI of mouse heads was performed as previously described^61,62^. After removal of the skin, ears, facial muscles, and mandibles, mouse heads were mounted in a modified 60-ml syringe using a custom plunger with three axially spaced levels, each holding four heads affixed with Skinaffix (Medline, Northfield, IL). The 60-ml syringe was chosen (instead of the typical 80-ml syringe) to accommodate the smaller head size of young mice. Polyethylene tubing filled with copper-doped water was inserted for orientation, and the syringe was filled with Fomblin to suppress background signal and improve shimming. Degassing was performed under vacuum for at least two hours before placing the syringe horizontally in the scanner. Imaging was conducted on a Bruker Maxwell 7T system (Avance Neo console, 170-mm bore, BGA-105S-HP gradients, 900 mT/m, 214 μs rise time, Paravision 360 version 3.6, Topspin 4.4.0) using a circularly polarized (CP) birdcage RF coil (40-mm inner diameter), which was well-matched to the syringe diameter to ensure optimal signal homogeneity. A motorized ATS bed enabled automated repositioning of three sample sets per session. A three-dimensional (3D) T2-weighted Fast-Spin Echo (FSE) sequence was employed for its superior gray/white matter contrast and cerebrospinal fluid (CSF) visualization, highlighting the ventricular system. Four heads were scanned simultaneously at a 125 µm isotropic resolution using the following parameters: Effective echo time (TEeff) = 42 ms, echo spacing (ES) = 7 ms, repetition time (TR) = 1,700 ms, acceleration factor (AF) = 16, number of averages (Nav) = 2, matrix size (Mx) = 205^3^, slice anti-aliasing (aa3) = 1.127, resulting in an Mx = 205 × 205 × 231. The field-of-view (FOV) was 25.6 mm^3^, with a bandwidth (BW) = 92.6 kHz (451.67 Hz/pixel), yielding a total imaging time of 2 hours and 48 minutes. Additionally, a 3D multi-gradient echo (MGE) sequence was acquired with six echo images under the same FOV and Mx using the following parameters: Nav = 4, TR = 45 ms, TE = 2.98 ms, ES = 3.68 ms, BW = 75.8 kHz (369.76 Hz/pixel), partial Fourier transform (pFT) in the phase direction of 1.33, and slice aa3 = 1.4, resulting in an effective Mx = 205 x 154 x 287, with a total scan time of 2 hours and 13 minutes.

### 3D brain segmentation and anatomical analysis

Post-acquisition, datasets were converted to NIfTI format for compatibility with ImageJ/Fiji^63^. The high resolution enabled precise 3D registration, anatomical identification, volume quantification, and rendering via Amira. The tissue contrast supported accurate segmentation and visualization of regions of interest throughout the whole brain.

### RT-qPCR

Total RNA was extracted using the RNeasy mini plus kit (QIAGEN, 74136). 2 µg of total RNA was used for reverse transcription using a SuperScript IV kit (Invitrogen, 18-091-050). SYBR Green Master Mix (Roche, 04887352001), 0.2 µl of cDNA and 0.25 µM primers were used in a 10 µl qPCR reaction. All RT-qPCR experiments were conducted with three technical replicates. Quantitative PCR was performed on a Roche LightCycler 480 instrument with standard qPCR program for SYBR Green Master Mix. Primers used in this study are listed in **Supplementary Table 1**.

### DNA copy number quantification by qPCR

mESC genomic DNA (gDNA) was extracted using the gDNA extraction kit (QIAGEN, 51306). Approximately 500 ng gDNA and 1 ng payload plasmid (containing a region of the mouse *Actb* gene on the backbone) were digested with restriction enzymes that do not recognize the PCR amplicons. Digested DNA was purified using 1x Sera-Mag Select beads (Cytiva, 29343052). 50 ng digested gDNA or 10 pg digested payload plasmid was used in a 10 µl SYBR Green qPCR reaction. Reference *Actb* primers (oWZ1683 and oWZ1684) and payload targeting primers (oRB394 and oRB395) were used for the qPCR. Payload DNA copy number is normalized to *Actb* and to the payload plasmid standard. A single-copy integrated payload has 1:2 payload:*Actb* ratio as *Actb* has two copies in mouse genome.

### Targeted sequencing

Targeted sequencing of genomic DNA was performed by following a previously described protocol^21^. Briefly, for each reaction, 500 ng of genomic DNA was used as input for sequencing libraries construction with the NEBNext Ultra II FS kit, selecting for fragments larger than 550 bp. Capture probes were prepared by combining mouse BAC DNA (RP23-334A11, BACPAC Resources Center), pSpCas9 plasmid, marker cassette 1 plasmid, and the XDP payload plasmid. The DNA mixture was labeled with Biotin-16-dUTP (Roche, 11431692103) using a nick translation kit (Sigma-Aldrich, 10976776001). The biotinylated probes were pre-hybridized and incubated with DNA libraries at 65 °C for 16–22 hours. Hybridized fragments were isolated using Streptavidin C1 beads (Invitrogen, 65002), followed by PCR amplification with the KAPA HiFi HotStart PCR kit (Roche, KK2602). After final purification, enriched libraries were sequenced on an Illumina NextSeq 550 platform using a 75-cycle kit. Sequencing data processing was performed as previously described^64^.

### 3’ RACE followed by Nanopore sequencing

For mESCs and human neural stem cells (NSCs), total RNA was extracted using the RNeasy mini plus extraction kit (QIAGEN, 74136). 2 µg of total RNA were used for reverse transcription with primer Qt (oWZ2245) following the SuperScript IV’s instructions. 1 µl cDNA was used as template, *TAF1* exon 30 forward primer 1 (oWZ2249) and primer Qo (oWZ2246) were used for the first round of PCR for 30 cycles. 1 µl of the first round PCR product was used as template, *TAF1* exon 32 forward primer (oWZ2250) and primer Qi (oWZ2247) were used for the second PCR round for 30 cycles. 3’ RACE PCR products were purified using the DNA Clean & Concentrator kit (Zymo Research, D4013). 100 ng amplicon was used as input for constructing Nanopore sequencing libraries by following the instructions of the Native Barcoding Kit 24 V14 (SQK-NBD114.24). For postmortem XDP human caudate nuclei, total RNA^44^ was reverse-transcribed using primer Qt (oWZ2245), following by first round of PCR of cDNA for 30 cycles using *TAF1* exon 30 forward primer 2 (oSC001) and Qo (oWZ2246), and second round of PCR for 30 cycles using *TAF1* exon 30 forward primer 3 (oSC002) and Qi (oWZ2247). A PCR barcoding kit (EXP-PBC-096) in combination with the Ligation Sequencing Kit (SLK-114) were used for library construction. Libraries were sequenced using a minION flowcell on a GridION sequencer. Reads were aligned using the minimap2/2.24 aligner to the custom hy*TAF1* genomic reference. Primer sequences are in **Supplementary Table 1**.

### Stranded RNA-seq and analysis

RNA integrity number (RIN) was determined by using a Bioanalyzer RNA nano kit (Agilent, 5067-1511). Samples with RIN>9 were subjected to library prep using a TruSeq Stranded Total RNA Ribo-Zero Gold kit (Illumina, RS-122-2303). Libraries were sequenced using a 100-cycles flowcell on a NovaSeq X+ sequencer. RNA-sequencing data were processed using the sns rna-star pipeline. Briefly, adapter sequences and low-quality bases were removed by using Trimmomatic (v0.36). The mouse reference genome was first modified by replacing the mouse *Taf1* genomic allele with the converted hy*TAF1* allele using the Reform web app (https://reform.bio.nyu.edu). Reads were aligned to the reformed reference using STAR (v2.7.3), guided by a modified Gene Transfer Format (GTF) file. Insert size means and standard deviations were calculated using Picard tools (v2.18.20). Gene-sample count matrices were generated with featureCounts (v1.6.3) and normalized by library size factors using DESeq2, followed by differential expression analysis. RPM-normalized BigWig files were created with deepTools (v3.1.0). Data were visualized using Integrative Genomics Viewer or plotted using GraphPad Prism 10.

### Growth rate measurement

A total of 5,000 cells were seeded per well in a gelatin-coated 96-well plate in 100 µl of ESM + 2i medium for expansion. After 3 days of culture at 37 °C in a humidified incubator with 5% CO_2_, cells were dissociated using 30 µl of Trypsin-EDTA at 37 °C for 5 minutes and quenched with 170 µl of ESM + 2i medium. The cell suspension was thoroughly dissociated by pipetting, and 90 µl of the resulting suspension was mixed with 10 µl of PrestoBlue reagent (Invitrogen, A13261) in a 96-well assay plate (Greiner Bio-One, 655892). Following 30 minutes of incubation at 37 °C, fluorescence was measured using a microplate reader (BioTek Synergy H1) with excitation at 560 nm and emission at 590 nm. Each mESC line was tested in four technical replicates.

### Immunoblotting

mESCs were harvested at 60–70% confluency. Cell number and viability were determined using a countess instrument. 1x RIPA buffer (Abcam, ab156034, diluted in deionized water) supplemented with 1x protease inhibitor cocktail (Roche, 11836170001) and Benzonase (1250 U/ml) was used for cells lysis (100 µl lysis per 1 million cells) for 30 minutes on ice. Cell lysates were supplemented with NuPAGE LDS Sample Buffer (Invitrogen, NP0007) and NuPAGE Sample Reducing Agent (Invitrogen, NP0004) and boiled at 80 °C for 10 minutes. Samples were centrifuged at 10000 x *g* for 1 minute at room temperature. Protein samples were loaded onto a 4–12% NuPAGE bis-tris midi gel for electrophoresis. Proteins were transferred to a PVDF membrane (EMD Millipore, IPFL00010) in NuPAGE transfer buffer (Invitrogen, NP0006, diluted in deionized water) supplemented with 10% methanol at 30 volts constant for 16 hours at 4 °C. PVDF membranes were incubated with blocking buffer (5% milk in TBST buffer) for 1 hour at room temperature, followed by incubation with primary antibody diluted in blocking buffer for 2 hours at room temperature. Primary antibodies used in this study: anti TAF1 (Abcam, ab188427 for Ab-A; Thermo Scientific, PA5-104490 for Ab-T), anti α-Tubulin (Sigma, T5168), anti PARP1 (Cell Signaling Technology, 9532), anti Cas9 (Cell Signaling Technology, 14697). After 5 washes in the TBST buffer, membranes were incubated with HRP-conjugated secondary antibody (Cell Signaling Technology, 7076 for anti-mouse IgG, 7074 for Anti-rabbit IgG) diluted in blocking buffer for 1 hour at room temperature. Signal was visualized using a chemiluminescent substrate (Thermo Scientific, A38555, 34075) in a ChemiDoc instrument (Bio-Rad).

### CUT&RUN

mESCs cultured on gelatin-coated plates were dissociated with 0.25% Trypsin-EDTA for 5 minutes followed by quenching with equal volume of ESM+2i medium. Cells were collected by centrifugation at 300 x *g* for 5 minutes at room temperature. Cell number and viability were determined using a countess instrument. CUTANATM CUT&RUN kit (EpiCypher,14-1048) was used for the following steps. In brief, 5×10^5^ cells were washed with Wash Buffer and pelleted at 600 x *g* for 5 minutes at room temperature. Cells were then incubated with activated Concanavalin A (ConA) beads at room temperature for 10[minutes. Antibodies against H3K4me3 and IgG were incubated with ConA beads-bound cells on a nutator at 4 °C overnight. Cells were permeabilized with 0.01% digitonin containing buffer. pAG-micrococcal nuclease was added to bind to the antibodies and then was activated with 2 mM CaCl_2_ for digestion for 2 hours at 4 °C. Digested DNA fragments were released by incubating the tubes at 37 °C for 10 minutes. DNA was purified by using 1.4xSPRI beads and eluted in 0.1xTE Buffer. NEBNext Ultra II DNA Library Prep Kit (E7645L) was used for sequencing library prep. Libraries were sequenced using a 75-cycle kit on an Illumina NextSeq 550 sequencer. Sequencing reads were aligned to the mouse genome (mm10) and a custom human *TAF1* genomic region separately.

### CRISPR activation

The VPR-dCas9 plasmid (#84247, pSLQ2814) and empty U6 sgRNA expression plasmid (#64046, pSB700) were from Addgene. Four gRNAs were cloned into pSB700 between two BsmBI sites using Golden Gate method. Four U6-sgRNA expression units were then amplified and inserted into the *Nhe*I site of the pSLQ2814 plasmid. Plasmids expressing both U6-sgRNA and VPR-dCas9 were delivered into converted-ΔSVA_D mESCs using the piggyBAC system (System Biosciences). After selecting the transfected cells with 1 µg/ml puromycin for two passages, cells were collected for total RNA extraction with RNeasy mini plus kit. RNA concentration was determined using a nanodrop instrument. One Step PrimeScript III RT-PCR kit (Takara, RR600B) was used in combination with FAM labeled probes (oWZ2487, oWZ2508) for hy*TAF1* mRNA, Cy5 labeled probe (oWZ1655) for *Actb* mRNA, as well as the corresponding primers for RT-qPCR. 50 ng total RNA was used for each RT-qPCR reaction.

## Acknowledgement

We thank the Collaborative Center for X-Linked Dystonia-Parkinsonism (CCXDP) and NIH (1RM1HG009491) for funding this work. We thank the core facilities at NYU Langone Health for their expertise. Specifically, members of the Rodent Genetic Engineering Laboratory (RRID:SCR_017925), members of the Preclinical Imaging (PCI) Laboratory (RRID:SCR_017937), members of the Genome Technology Center (RRID: SCR_017929), and members of the Applied Bioinformatics Laboratories (RRID:SCR_019178), which are partially supported by the Cancer Center Support Grant P30CA016087 at NYU Langone Health Laura and Isaac Perlmutter Cancer Center. Part of the work conducted at the PCI laboratory was supported in part by the NIH Shared Instrumentation Grant 1S10OD018337-01 and the NIBIB Biomedical Technology Resource Center Grant NIH P41 EB017183. We thank Dr. Yubao Wang for kindly sharing the dTAG constructs and the dTAG degrader. We thank Emily Huang, Hannah J. Ashe and Chloe Palumbo for assisting with targeted sequencing. We also acknowledge the Belfer Neurodegeneration Consortium and the Carol and Gene Ludwig Family Foundation (S.A.L.). We thank the Leon Levy Foundation for postdoctoral fellowship in neuroscience award (P.P.).

## Author contributions

J.D.B. conceived the study. W.Z., R.B. designed the experiments. Y.Z., K.B., G.E. assembled DNA constructs. W.Z., H.L.A. engineered and characterized mESCs. S.Y.K. and W.Z. established the mouse colonies. P.P., K.C.L. performed the nestin-Cre mouse crossing, brain dissection for genomic DNA and total RNA extraction. W.Z., S.C., A.M.W., C.V. performed 3’ RACE, sequencing and data analysis. R.B., M.T.M. supervised targeted genome sequencing and data analysis. W.Z. performed RNA sequencing, Q.J. performed GSEA and motif analyses. W.Z. performed RT-qPCR and immunoblotting experiments. Y.Z.W. devised the thigh-throughput *ex vivo* MRI strategy. S.M. developed the MRI pulse sequence protocol with assistance from N.R. and O.M. during preliminary testing. N.R. carried out all sample preparations for MRI scanning, image processing, and post-analysis. O.M. supported image acquisition. J.D.B., R.B., S.A.L., H.T.M.T., D.C.B. supervised the study. W.Z. drafted the manuscript, J.D.B., R.B. edited the manuscript and all authors reviewed the manuscript.

## Supplementary Tables

**Supplementary Table 1.**
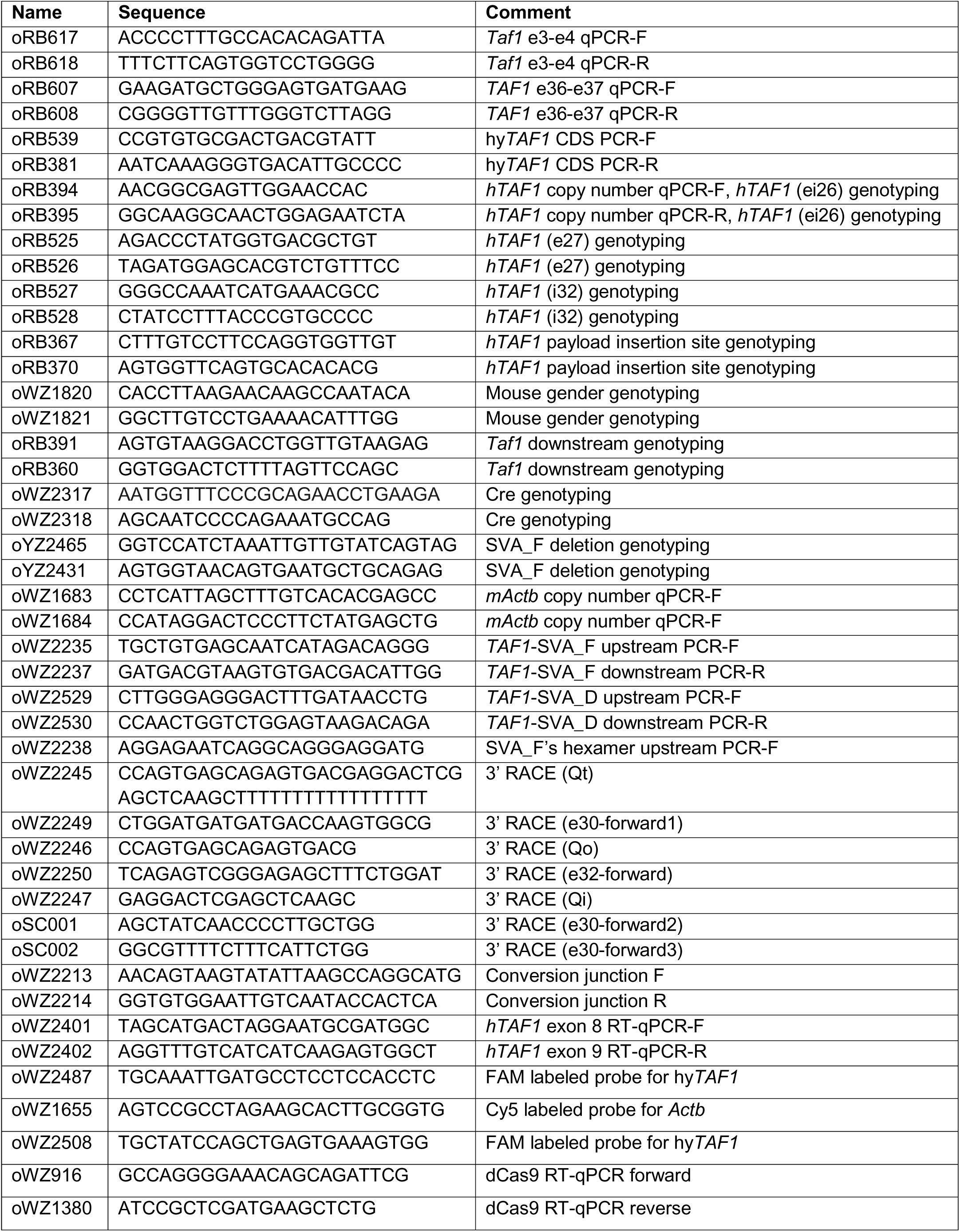
Primers used in this study.

**Supplementary Table 2.**
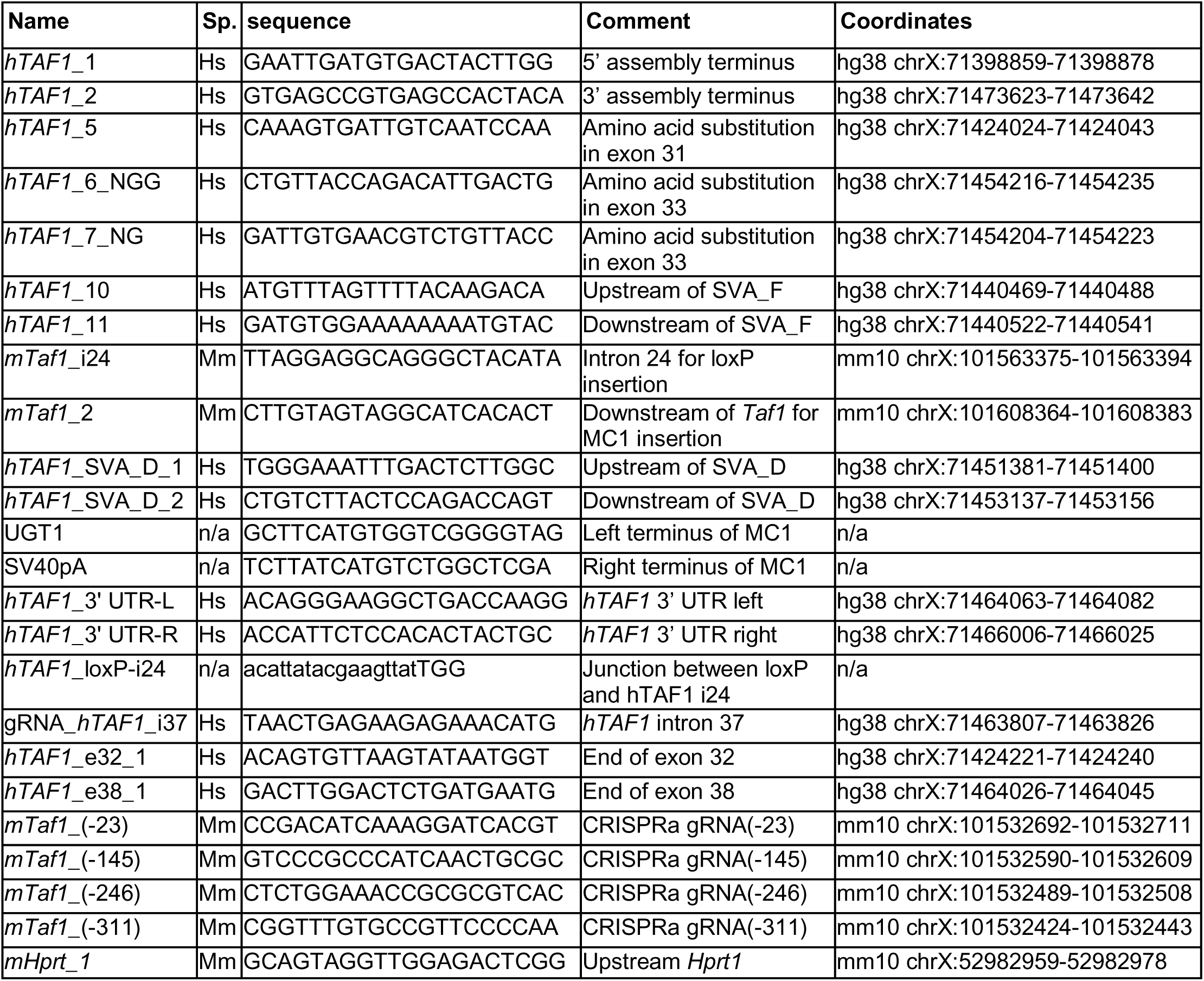
sgRNAs used in this study.

---

**Figure. S1.**
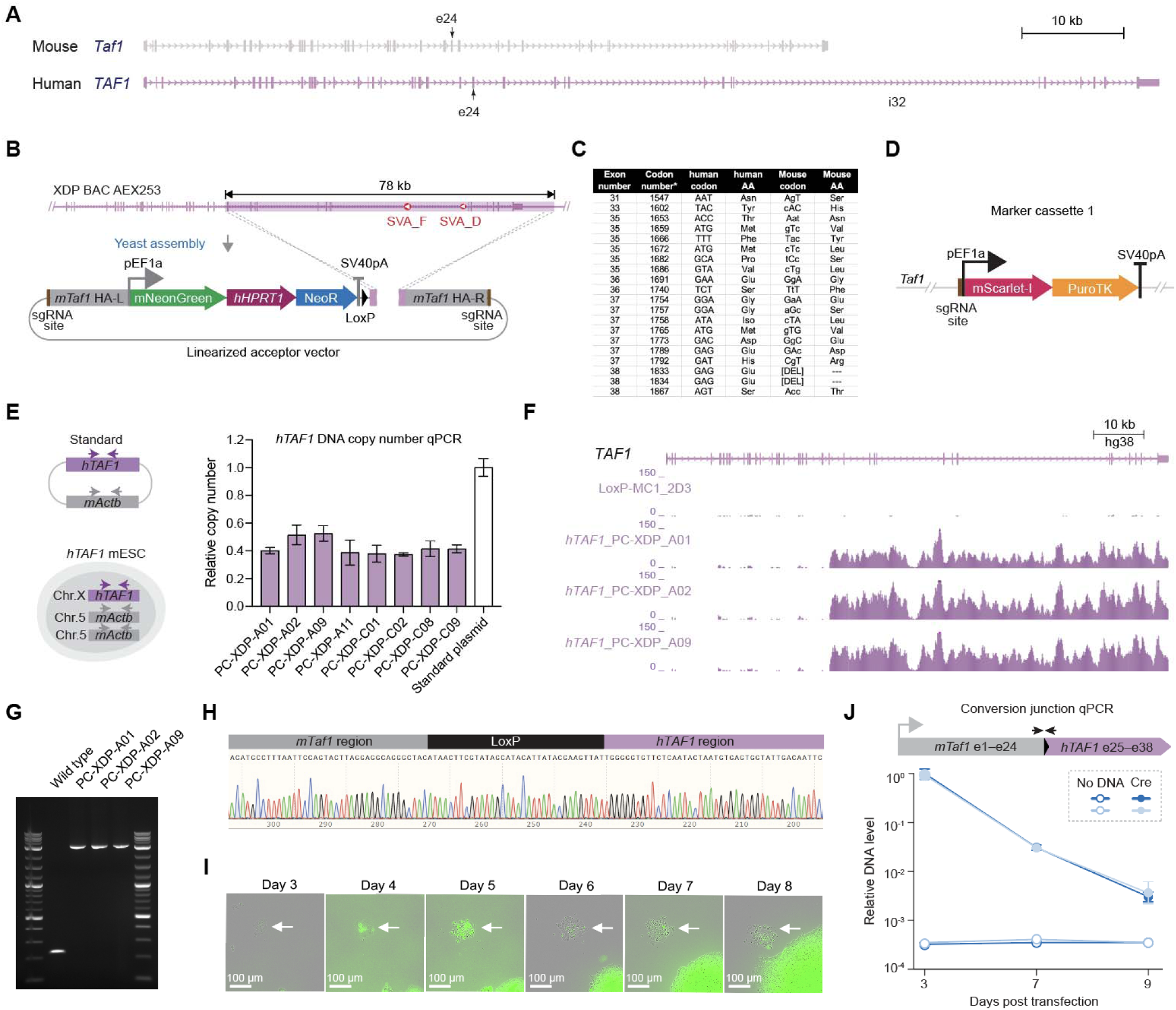
Engineering and validation of *TAF1* humanized mESCs. Related to Figure 1. (A) Schematics of mouse *Taf1* gene (grey) and human *TAF1* gene (purple). Tall bars are exons; short bars on both ends are untranslated regions (UTRs); regions marked with arrows are introns. Relative gene lengths are drawn to scale. Exon 24 (e24) and intron 32 (i32) are denoted. (B) Cloning strategy for inserting human *TAF1* exon 25 through the 3′ UTR into a yeast acceptor vector compatible with mESCs genome engineering. The human *TAF1* region of interest was released from XDP BAC AEX253 (see Methods) by *in vitro* Cas9 digestion. The acceptor vector was linearized by *Sal*I digestion, and two pre-installed 200 bp homology arms (purple boxes) mediated recombination with the termini of the human *TAF1* fragment in yeast. (C) Substituted amino acids and their codons. In the mouse codon column, lowercase letters indicate variants introduced to the human TAF1 sequence to induce substitutions. Asterisk, codon number refers to NM_004606.5 *Homo sapiens* TATA-box binding protein associated factor 1, transcript variant 1. (D) Structure of marker cassette 1 (MC1). (E) Quantification of human *TAF1* DNA copy number in engineered mESC clones^18^. A plasmid containing both the human *TAF1* payload and a fragment of mouse *Actb* (1:1 ratio) was used as a standard. Values were normalized to *Actb* and then to the standard. Bars represent mean ± SD of three technical replicates. A relative copy number of 0.5 supports single-copy integration compared with the two copies of mouse endogenous *Actb*. (F) Targeted sequencing of engineered mESC clones. Reads were aligned to a human genomic reference, hg38. A parental mESC clone with loxP and MC1 insertions was used as a control. Three independent pre-converted XDP mESC clones were included. (G) PCR detection of SVA_F (∼3 kb) in three engineered XDP mESC clones. Human colorectal carcinoma cell line HCT116 genomic DNA was used as wild-type human genomic DNA control. Ladders, 1 kb Plus DNA Ladder (NEB). (H) Sanger sequencing of the conversion junction in transiently Cre-transfected (2-day post transfection) XDP mESCs, confirming conversion fidelity. (I) Live-cell imaging of XDP mESCs post conversion. Arrows indicate a transiently converted XDP clone exhibiting signs of apoptosis that could not form a viable colony; GFP positive colonies grow into large colonies that remain unconverted. Scale bar is 100 μm. (J) Time-course quantification of conversion junction DNA abundance in XDP mESCs. A genomic mouse *Actb* primer pair (oWZ1683 and oWZ1684) was used as an internal reference. Cells transfected with Tris-EDTA (TE) buffer served as negative control.

**Figure. S2.**
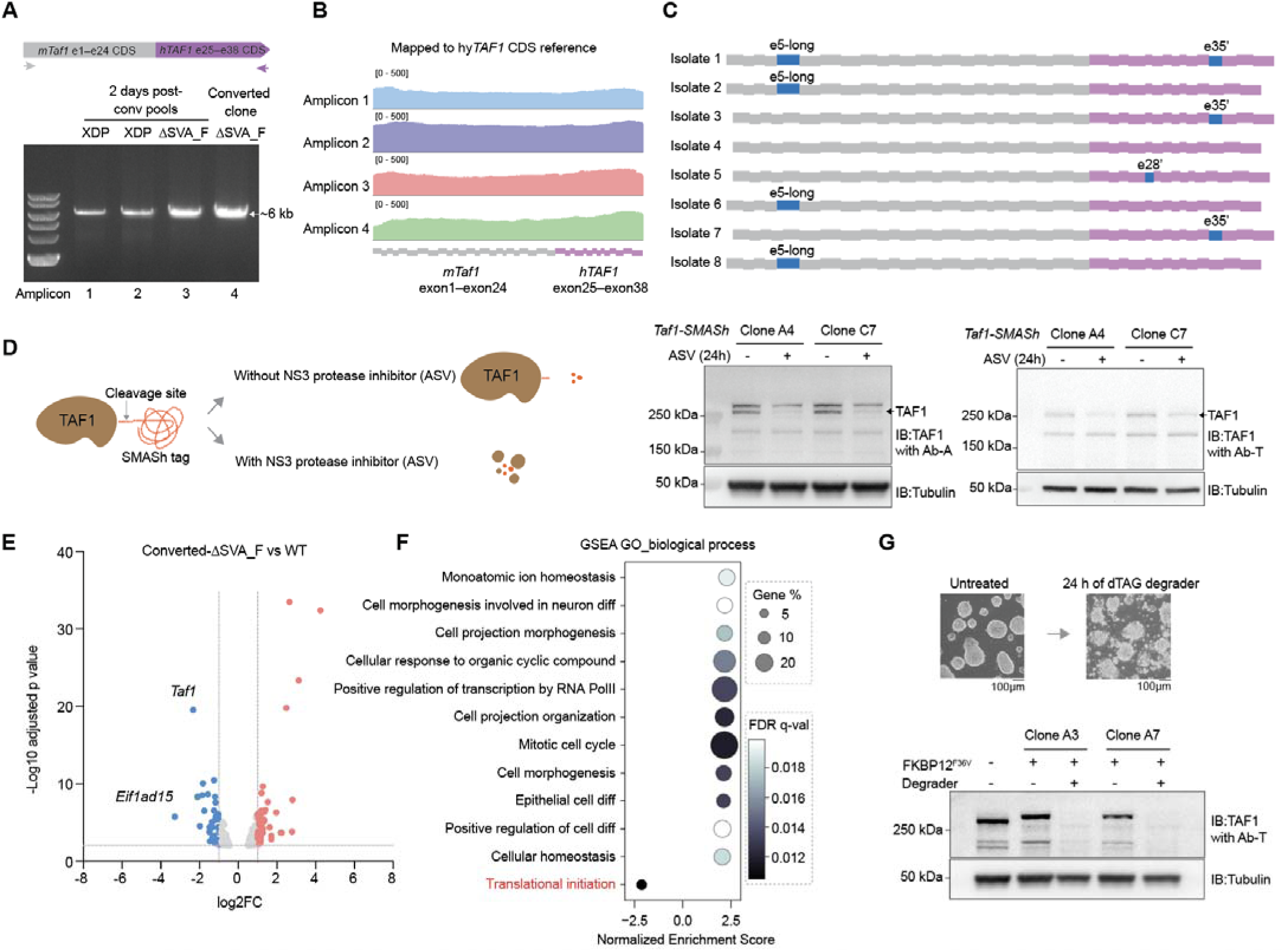
Validation of hy*TAF1* transcripts and TAF1 depletion. Related to Figure 2. (A) PCR amplification of full-length hy*TAF1* cDNA. Grey arrow marks the mouse *Taf1* forward primer (oRB539), purple arrow marks the human *TAF1* reverse primer (oRB381). RNA was extracted from a pool of cells collected two days post Cre transfection or single colony-isolated converted-ΔSVA_F cells, followed by first strand cDNA synthesis. PCR products were resolved on a 1% agarose gel. Ladder, 1 kb plus DNA ladder from NEB. (B) Long-read sequencing of the amplicons in panel (a). Reads were aligned to a custom hy*TAF1* coding sequence reference. Grey and purple rectangles below indicate mouse and human exons, respectively. (C) TOPO cloning and sequencing of eight clonal isolates from the amplicons in panel (a). Alternative exons are highlighted in blue; e5-long: mm10 chrX:101540760–101541001; e28’: hg38 chrX:71421145–71421225; e35’: hg38 chrX:71459150–71459251. Grey and purple rectangles denote mouse and human exons, respectively. (D) TAF1 antibody validation using the SMASh degron system^31^. SMASh degron cleaves itself off in the absence of a protease inhibitor, producing minimally tagged protein (6 aa) with nearly unchanged molecular weight. The tagged protein is degraded after blocking the protease activity (left panel). Wild-type mESCs engineered with a TAF1 C-terminal SMASh tag were treated with 1 μM asunaprevir (ASV) for 24 hours. TAF1 levels were assessed by western blotting using two commercial antibodies (Ab-A and Ab-T, see Methods). Arrows indicate TAF1 bands. α-Tubulin served as loading control. (E) Volcano plot of differentially expressed genes between converted-ΔSVA_F and wild-type mESCs. Fold change cutoff is 2, adjusted p-value cutoff is 0.01. Red and blue dots represent significantly up- and downregulated genes in converted-ΔSVA_F cells, respectively. (F) Gene set enrichment analysis (GSEA) using the Gene Ontology Biological Process subset for mouse. Pathways with FDR q-value < 0.02 are shown. (G) Brightfield images of mESC morphology before and after 24 hours of dTAG degrader treatment (upper panel). Western blot analysis showing near-complete depletion of TAF1 in degrader-treated mESCs (bottom panel). Two independent dTAG clones were analyzed. Wild-type mESCs were used as negative control. α-Tubulin served as loading control.

**Figure. S3.**
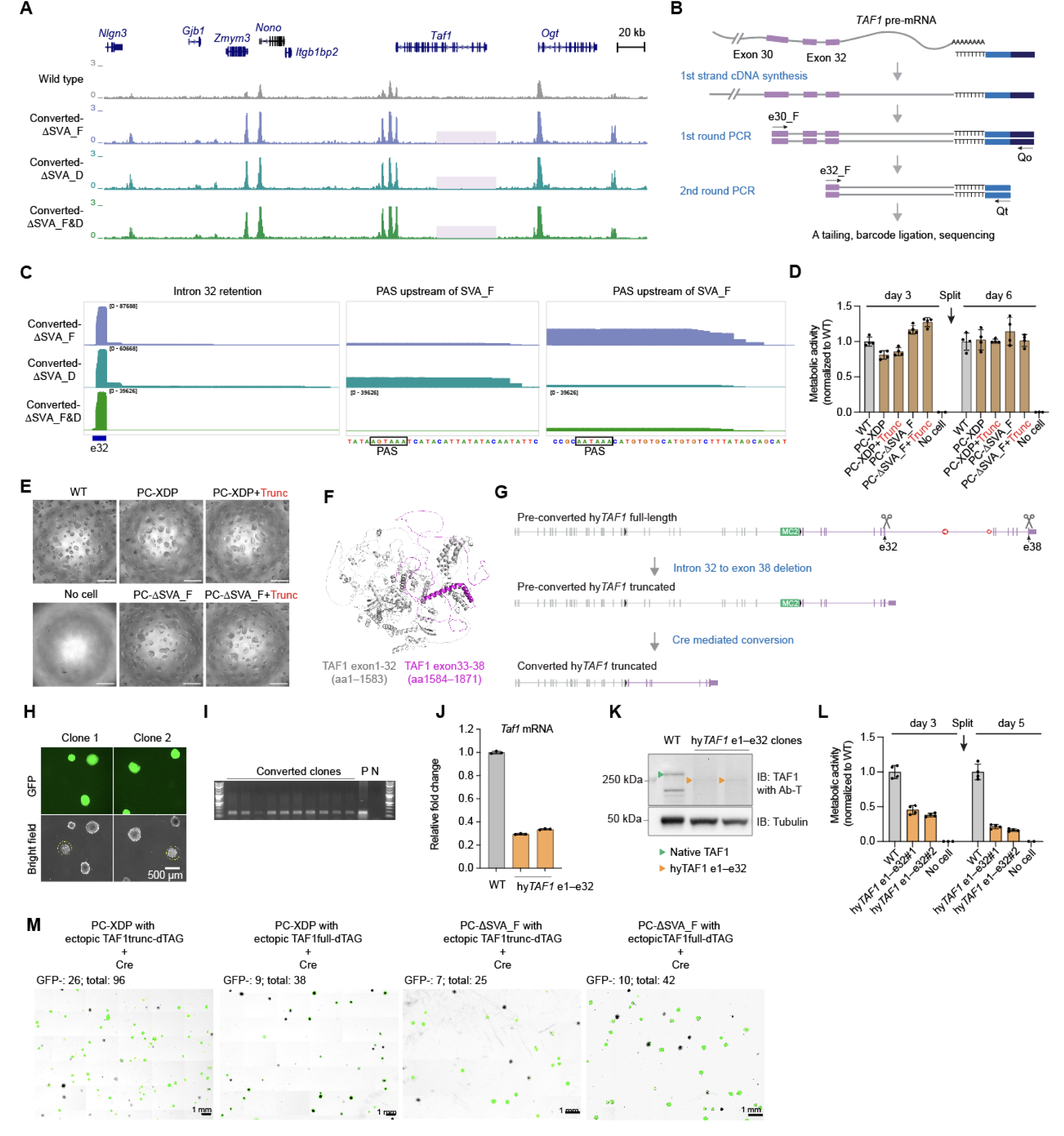
Characterization of transcriptional interference and functional testing of truncated TAF1. Related to Figure 3. (A) mESCs H3K4me3 CUT&RUN. Sequencing reads were mapped to the mouse genome. Transparent purple boxes mark the region of mouse *Taf1* that is excised following Cre-mediated conversion. (B) Schematic of 3’ RACE protocol. An oligo-dT-containing reverse primer (oWZ2245) was used for first strand cDNA synthesis. Nested PCR was used to capture the prematurely terminated transcripts. 3’ RACE products were sequenced by nanopore long-read sequencing on a GridION instrument. (C) Zoomed-in views of the 3’ RACE sequencing data from Fig. 3D, highlighting representative peaks: one at the beginning of intron 32 (left), one upstream of SVA_F (middle), and one downstream of SVA_F (right). PAS, polyadenylation signal. (D) Comparison of metabolic activity using resazurin, a cell-permeable redox indicator, which is metabolized by viable cells, producing a fluorescent signal proportional to cell number. Bars represent mean ± SD from four technical replicates. (E) Bright field views of mESC growth on gelatin-coated 96-well plate. Scale bar is 500 µm. (F) Predicted structure of the mouse TAF1 protein (AlphaFold Protein Structure Database). Grey and magenta regions correspond to exons 1–32 and 33–38, respectively. (G) Experimental design to assess the functionality of the truncated hy*TAF1* (exons 1–32) allele. (H) Representative fluorescence images showing successfully converted hy*TAF1* exon 1–32 colonies (yellow dashed circles, GFP-negative) post Cre transfection. (I) PCR genotyping of the conversion junction in converted hy*TAF1* exon 1–32 clones. P, positive control. N, negative control. (J) Quantification of hy*TAF1* mRNA levels in converted hy*TAF1* exon 1–32 clones. Bars represent mean ± SD of three technical replicates. (K) TAF1 western blot analysis of converted hy*TAF1* exon 1–32 mESC clones. α-Tubulin serves as loading control. (L) Comparison of metabolic activity. Bars represent mean ± SD from four technical replicates. (M) Raw images showing single colonies of Cre transfected pre-converted-XDP mESCs with ectopic expression of truncated TAF1-dTAG or full-length TAF1-dTAG; and Cre transfected pre-converted-ΔSVA_F mESCs with ectopic expression of truncated TAF1-dTAG or full-length TAF1-dTAG. Images were acquired 7 days post Cre transfection using a CellCelector instrument.

**Figure. S4.**
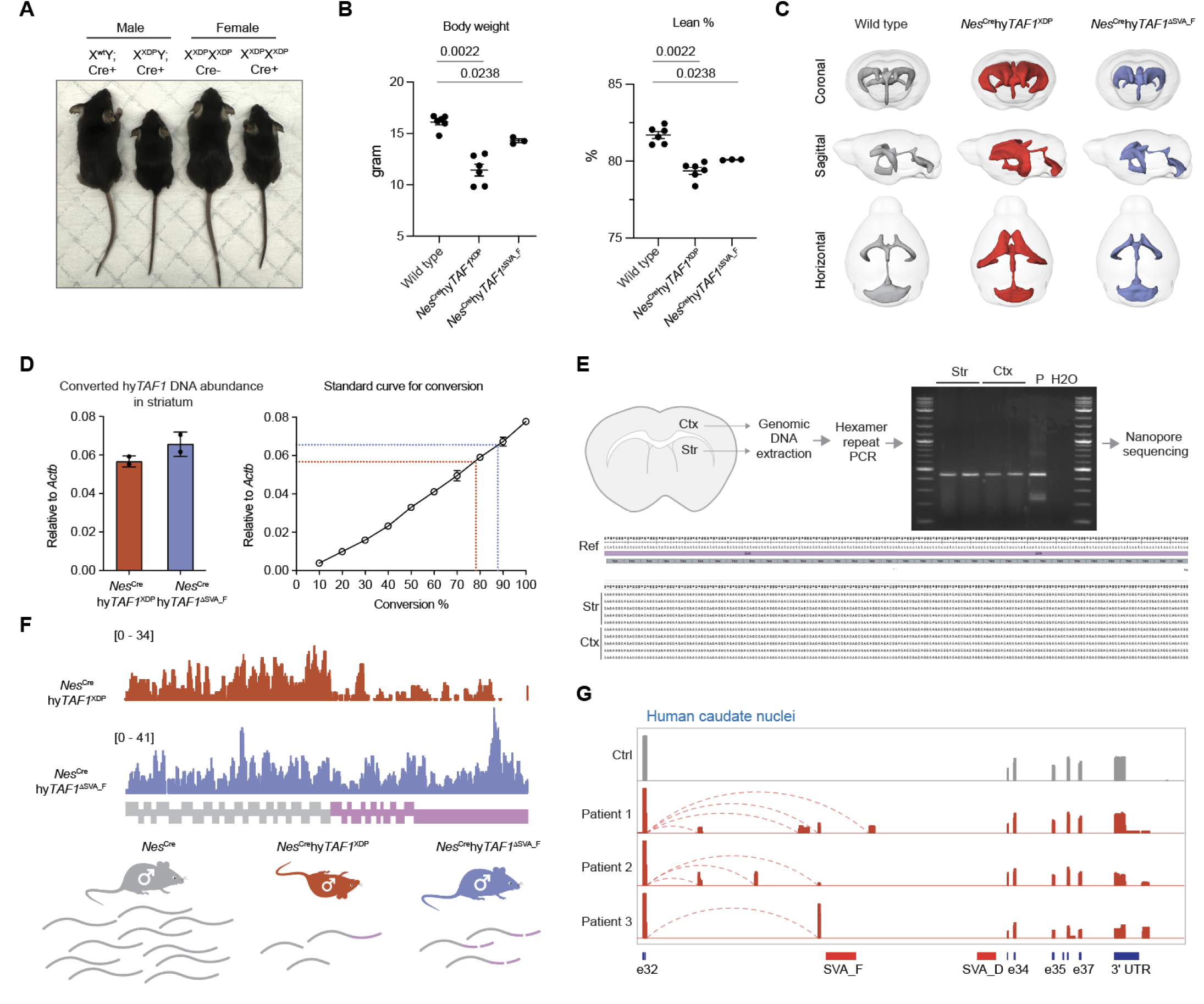
XDP mice and hy*TAF1* mRNA characterization. Related to Figure 4. (A) Representative mice image at 3 weeks of age. The four mice were littermate progeny from *Nes*^Cre+/-^ hy*TAF1*^XDP/WT^ female and *Nes*^Cre-/-^hy*TAF1*^XDP/Y^ cross. (B) Body weights and lean percentages of wild-type (n=6), *Nes*^Cre^hy*TAF1*^XDP^ (n=6) and *Nes*^Cre^hy*TAF1*^ΔSVA_F^ (n=3) male mice. Bars represent mean ± SEM. Two-tailed Mann-Whitney *U* test. (C) 3D rendering of ventricular systems from the MRI scans in Fig. 4D. (D) Quantification of *in vivo* conversion efficiency in *Nes*^Cre^hy*TAF1*^XDP^ and *Nes*^Cre^hy*TAF1*^ΔSVA_F^ striata (left). A standard curve (right) was established by qPCR using known ratios of converted to non-converted genomic DNA, enabling estimation of the proportion of converted DNA in brain tissues. (E) Analysis of the hexamer repeat length within the SVA_F in cortex and striatum of 3-week-old converted XDP mice. The repeat-containing region was PCR-amplified from genomic DNA and resolved on a 1% agarose gel (top). Then the PCR-amplicons were TOPO-cloned, individual bacterial isolated were sequenced by nanopore sequencing. Five independent TOPO clones were analyzed (bottom). (F) RNA-seq reads from conditionally converted striata mapped to the custom hy*TAF1* coding sequence (top). Schematic model (bottom) illustrating that reduced functional hy*TAF1* transcript in the brain underlines the postnatal lethality observed in *Nes*^Cre^hy*TAF1*^XDP^ mice. (G) 3′ RACE sequencing of *TAF1* transcripts in caudate nucleus from three XDP patients and one unaffected control, revealing premature termination events associated with the SVA_F insertion. Dashed arches indicate contiguous reads spanning exon 32 and the cryptic exons.

**Figure S5.**
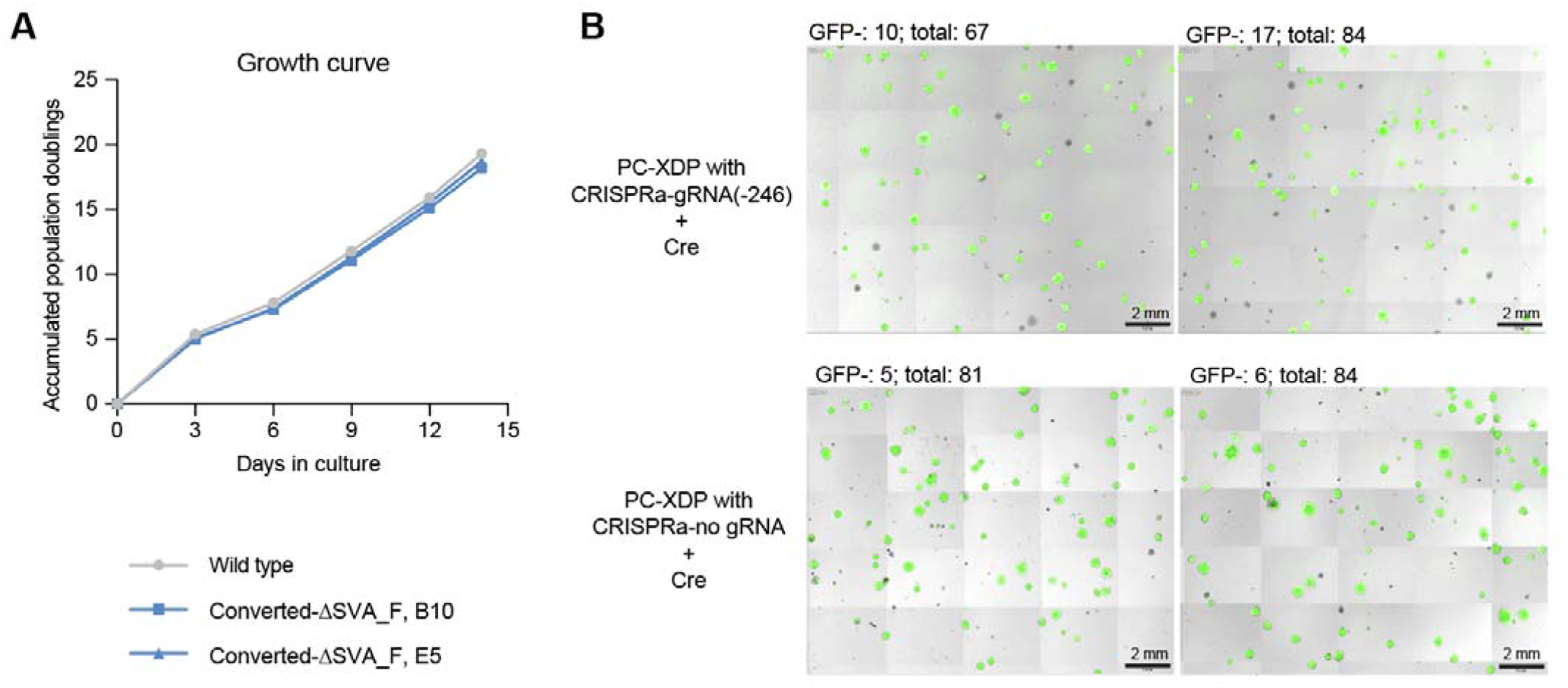
CRISPRa assisted conversion. Related to Figure 5. (A) Growth rate comparison between wild type mESCs and two clones of converted-ΔSVA_F mESCs. Population doublings were computed by counting mESCs number changes for 5 passages in 14 days. (B) Raw images showing single colonies of Cre transfected pre-converted-XDP mESCs with CRISPRa-gRNA (−246) integrated (up) or CRISPRa-no gRNA integrated (bottom). Images were acquired 7 days post Cre transfection using a CellCelector instrument.

